# A new GABAergic projection from the BNST onto accumbal parvalbumin neurons controls anxiety

**DOI:** 10.1101/767228

**Authors:** Qian Xiao, Xinyi Zhou, Pengfei Wei, Li Xie, Yaning Han, Bifeng Wu, Jie Wang, Aoling Cai, Fuqiang Xu, Yi Lu, Jie Tu, Liping Wang

**Affiliations:** Shenzhen Key Lab of Neuropsychiatric Modulation, Guangdong Provincial Key Laboratory of Brain Connectome and Behavior, the Brain Cognition and Brain Disease Institute (BCBDI), Shenzhen Institutes of Advanced Technology, Chinese Academy of Sciences (CAS); Shenzhen-Hong Kong Institute of Brain Science-Shenzhen Fundamental Research Institutions, Shenzhen, 518055, China; University of Chinese Academy of Sciences, Beijing100049, P.R. China; Department of information technology and electrical engineering, ETH Zurich; State Key Laboratory of Magnetic Resonance and Atomic and Molecular Physics, Wuhan Institute of Physics and Mathematics, CAS center for Excellence in Brain Science and Intelligence Technology, Chinese Academy of Sciences, Wuhan 430071, P.R. China; Wuhan National Laboratory for Optoelectronics, Huazhong University of Science and Technology, Wuhan 430074, P.R. China

## Abstract

The prevailing view is that parvalbumin (PV) interneurons play modulatory roles in emotional response through local medium spiny projection neurons (MSNs). Here, we show that PV activity within the nucleus accumbens shell (sNAc) is required for producing anxiety-related avoidance when mice are under anxiogenic situations; sNAc^PV^ neurons exhibited high excitability in chronically stressed mice model, which generated excessive maladaptive avoidance behavior in an anxiogenic context. We also discovered a novel GABAergic projections from the anterior dorsal bed nuclei of stria terminalis (adBNST) to sNAc^PV^ neurons; optogenetic activation of these afferent terminals in sNAc produced an anxiolytic effect via GABA transmission. Next, we further demonstrated that chronic stressors attenuated the inhibitory synaptic transmission at adBNST^GABA^ 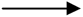 sNAc^PV^ synapses, which in turn explains the hyperexcitability of sNAc^PV^ neurons on stressed models; therefore, activation of these GABAergic afferents in sNAc rescued the excessive avoidance behavior related to anxious state.

Our findings reveal the coordination between BNST and NAc through an inhibitory architecture in controlling of anxiety-related response and provide a neurobiological basis for therapeutic interventions in pathological anxiety.

## Introduction

Stressors and stress responses are critical for guiding both approach and avoidance behaviors in animals and humans. Exposure to chronic, unpredictable stressors leads to increased anxiety responses, including excessive avoidance behavior, and this exposure has been adopted to study anxious state-related behaviors^1, 2^. The bed nucleus of the stria terminalis (BNST), a subregion of the extended amygdala, is a critical node in the stress response^3, 4^. Recent work on human drug addiction has also demonstrated a role of BNST in withdrawal-related anxiety and relapse, indicating an intrinsic link between this stress response region and the reward system. The nucleus accumbens (NAc) is a vital component in the reward circuitry^5–7^, which responds to stress signals^8, 9^ and has a dominant effect on anxiety regulation^10^. However, with the exception of one 15-year-old anatomical observation^11^, the connectivity between NAc and BNST, and whether it is necessary in producing anxiety-related behavior, remains unexplored. Furthermore, GABAergic efferents originating from the BNST are predominantly sent downstream^12, 13^; however, the nature and function of any GABAergic input to NAc is unknown. Medium spiny neurons that express dopamine 1 and 2 receptors (D1- and D2-MSNs) are the predominant neural population in the NAc. Regarding anxiety regulation, it is accepted that D1-MSNs are not involved in anxiety-related behavior, but play roles in modulating reward-related responses^14, 15^ whereas D2-MSNs may regulate anxiety-related aversion or avoidance behavior^16^. However, the role of D2-MSNs are not entirely clear because other work points to a role in reward seeking, but not anxiety-related behavior^17, 18^. Based on these very different findings, we predicted that there is another neuronal type within the NAc that contributes to anxiety-related behavior. One possibility is PV GABAergic interneurons, which comprise only 3-4% of all neurons in the NAc^19, 20^. In other brain regions, these neuron regulate fear response^21^, anxiolysis^22^, alcohol addiction^23^, and reward seeking^3, 24^. However, we know less about the function of accumbal PV neurons and the inputs they receive, and nothing about any possible role in anxiety related behavior.

We addressed these important questions regarding the neural mechanisms underlying the expression of avoidance of anxiogenic stimuli in both healthy and pathological anxiety models. Combining functional MRI signaling, GCaMP-based fiber photometry recording, genetically modified virus tracing and both optogenetic and chemogenetic neuronal manipulations, we show that in an anxious state, functional connectivity between BNST and NAc was increased; PV neurons within the NAc shell (sNAc) exhibited high excitability in chronically stressed mouse model that displayed excessive maladaptive avoidance during anxiogenic stimuli; and further, activation of these accumbal PV neurons promoted an avoidance coping response in healthy mice. A new GABAergic afferent from the anterior dorsal BNST (adBNST) was uncovered, which directly innervates sNAc^PV^ neurons. Optogenetic activation of these GABAergic terminals in sNAc produced an anxiolytic effect, which was mediated by sNAc^PV^ cells; activation of these inhibitory inputs from adBNST to sNAc rescued the excessively anxious state of the stressed mice.

Therefore, our results reveal a previously undescribed circuit mechanism, defined by neuronal type, that shapes the coordination between BNST^GABA^ and NAc^PV^ cells in response to anxiogenic stimuli under both physiological and pathological conditions.

## Materials and methods

### Mouse Models

All experiments were approved by the Shenzhen Institutes of Advanced Technology, Chinese Academy of Sciences Research Committee, and all experimental procedures involving animals were carried out in strict accordance with the Research Committee’s animal use guidelines. Surgery was performed under full anesthesia, and every effort was made to minimize animal suffering. Male and female mice (6-14 weeks) were used in this study. We used the following mouse lines : PV-Cre (B6;129P2-Pvalbtm1(cre)Arbr/J, Jackson Laboratory, stock No.008069), D1-Cre (B6.Cg-Tg(Drd1a-cre)262Gsat/Mmcd, MMRRC, 030989-UCD), D2-Cre (B6.FVB(Cg)-Tg(Drd2-cre)ER44Gsat/Mmcd, MMRRC,032108-UCD),. C57BL/6J wild-type mice were also used.

### Behavioral tests

All behavioral tests were performed blind to mice genotypes. Groups of mice were age-matched (8-14 weeks). All mice were handled for 15-30 min per day for three days before behavioral assays to reduce stress introduced by contact with experimenter.

#### 1) Elevated plus maze test

A plastic elevated plus maze consisting of a central platform (5×5 cm) with two white open arms (25×5×25 cm) and two white closed arms (25×5×25 cm) extending from it in a plus shape was used. The maze was elevated 65 cm above the floor. Mice were individually placed in the center with their heads facing a closed arm. The number of entries and the amount of time spent in each arm type were recorded.

#### 2) Open-field test

A plastic open field chamber (50×50 cm) was used and conceptually divided into a central field (25×25 cm) and a peripheral field for analysis. Each mouse was placed in the peripheral field at the start of each test. The open field test consisted of a 10 min session and mice locations were monitored/tracked using Anymaze software.

#### 3) Unpredictable chronic mild stress procedure (UCMS)

The UCMS protocol was performed as previously described ^51, 52^ with modification. Mice were exposed to environmental stressors for three weeks. One of the following stressors were presented during each daily session in a random order over three weeks: (i) restraint, where each mouse was placed in a tube (50 mL) for two hours without access to food or water, (ii) a wet environment where water was added (such that bedding was damp but not overly wet) to a housing cage containing mice for twelve hour sessions, and (iii) squeezing, where four mice were housed into a box (3 × 5 ×7 cm) for two hours, without access to food or water.

### Fiber photometry

Fiber photometry allows for real-time excitation and recording of fluorescence from genetic encoded calcium indicators in freely moving mice. Mice were habituated to the fiber patch cord for at least 15 min per day for three days before tests were conducted inside home cages. The fiber photometry system (ThinkerTech, Nanjing, China) consisted of an excitation LED light (480 nm; CREE XPE), reflected off a dichroic mirror with a 435-488 nm reflection band and a 502-730 nm transmission band (Edmund, Inc), coupled into a 200 μm 0.37 NA optical fiber (Thorlabs, Inc) by an objective lens. The laser intensity at the fiber tip was approximately 25-30 µW. GCaMP6_m_ fluorescence was recorded using the same objective, transmitted by the dichroic mirror filtered through a green fluorescence protein (GFP) bandpass emission filter (Thorlabs, Inc. Filter 525/39), and detected by the sensor of an CMOS camera (Thorlabs, Inc. DCC3240M).

A Labview program was developed to control the CMOS camera which recorded calcium signals at 50 Hz. Behavioral event signals were recorded by a DAQ card (NI, usb-6001) at 1000 Hz using the same program.

### Single-unit and local field potential (LFP) recordings in freely-moving mice

Both naive and chronically stressed mice (aged 8-12 wk) were anesthetized with isoflurane (4.0% for induction and set-up on the animal bed, 0.8%-1% during experiments) in a 20% O_2_/80% air mixture). Body temperature was maintained at 36-37 °C with a heating pad. For single-unit recording, mice were secured in a stereotaxic apparatus and a custom-made screw-driven microdrive containing eight tetrodes (four wires wound together) was unilaterally implanted in the left NAc shell; and for LFP recordings, custom-made stereotrodes (two wires wound together) were unilaterally implanted in both left NAc shell and aBNST. Each stereotrode was housed in a silica tube and consisted of two individually insulated platinum-iridium wires (17 µm inner diameter). The electrodes were modified by electrochemical deposition of platinum to reduce their impedance to ∼500 KΩ. The skull was leveled using bregma and lambda landmarks, and screws were implanted on the anterior and posterior portions of the skull to serve as reference.

Coordinates were measured from bregma and depth was calculated from the brain surface. The electrodes were implanted through burr holes in the skull aimed at the following coordinates: AP 1.35 mm, ML 1.35 mm, and DV -4.85 mm for NAc shell and AP 0.20 mm, ML 0.80 mm, and DV -4.05 mm for aBNST. The microdrive electrode was attached to a micromanipulator and moved gradually to a position about 400 μm above the desired depth. The electrodes were anchored to the microdrive that made it possible to advance along the dorsal-ventral coordinates. Following surgery, mice were allowed to recover for at least one week and were then habituated to experimenter handling. During recording, electrodes were connected to a unitary gain headstage (Plexon, Dallas, TX) connected to a 32-channel preamplifier (Plexon, Dallas, TX). Once mice were familiar with the recording setup they were connected to the headstage preamplifier in their home cages for two daily sessions of 20 min each. Neurophysiological signals were recorded with a 64-channel Multichannel Acquisition Processor (Plexon, Dallas, TX) and mouse positions were tracked using an overhead camera (30 Hz). Wideband signals were recorded at 40 kHz and LFP signals were acquired at 1 kHz.

At the end of each experiment, each mouse was deeply anesthetized with 10% chloral hydrate (0.4 mg/kg) and transcardially perfused with PBS, then 4% paraformaldehyde (PFA) (wt/vol). Brains were dissected and postfixed at 4 °C in 4% PFA overnight. Brains were then frozen and cut into 40 μm coronal slices and mounted on slides. Recording sites were marked with electrolytic lesions prior to perfusion and examined under a microscope to confirm recording locations.

### Resting state MRI

All mice were initially anesthetized with isoflurane (4.0% for induction and set-up on the animal bed, 0.8%-1% during experiments) in a 20% O_2_/80% air mixture. Body temperature was maintained at 37 °C using warm water circuitry. MRI experiments were conducted using a Bruker Biospec70/20USR small animal MR system (Bruker BioSpin MRI, Ettlingen, Germany) operating at 300 MHz (7T). Breathing rate, heart rate and blood oxygen saturation were monitored using a pulse oximeter positioned at the hind limb and a pressure-sensitive sensor under the mouse chest (MR-compatible Small Animal Monitoring & Gating System, SA Instruments, Inc.).

A planar receive-only surface coil with 20 mm diameter was used in combination with a detunable partial volume transmit coil (BrukerBioSpin MRI, Ettlingen, Germany). T2 anatomical reference scans in the coronal plane were acquired using a spin echo (Turbo-RARE) sequence: field of view (FOV) = 18×18 mm^2^, matrix dimension (MD) = 256×256, repetition time (TR)= 5000 ms, echo time (TE) = 12 ms, RARE factor = 8, number of averages (NA) = 4, spatial resolution = 0.0703125×0.0703125×0.8 mm^3^, 12 slices, no gap.

Resting state data sets were then acquired using single shot gradient echo EPI (Echo Planar Imaging) with TR 1000 ms and TE 16 ms. Twenty coronal slices (using the same procedure as the T2 anatomical images above) were recorded with a FOV of 1.8×1.8 mm^2^ and matrix size of 64×64, resulting in voxel dimensions of 0.28125×0.28125×0.8 mm^3^. Each resting state fMRI dataset comprised 300 repetitions, resulting in a scanning time of 10 min 40 s each. The bandwidth used was 200 kHz (6250 Hz/voxel). Preprocessing was done using SPM12 (http://www.fil.ion.ucl.ac.uk/spm/software/spm12/) to eliminate head movement and image shift by co-registering with the anatomical image, and then Gaussian smoothing was performed for every slice to improve the signal to noise ratio. To estimate functional connectivity, 4-5 voxels in each bilateral NAc image were selected as ROI (seed points), using REST (http://restfmri.net/forum/index.php) and home-written algorithms using Matlab2014a (www.mathworks.com).

### Stereotactic virus injection and optogenetic manipulation

Cre-mice (6-12 weeks) were used for stereotactic viral injections in the NAc shell. During isoflurane anesthesia (as above), the skull was exposed via small incision and a small hole was drilled for injection. A modified microliter syringe (Hamilton) with a 22-gauge needle was used: the tip of the needle was placed at the target region and the injection was performed at a speed of 100 nl/min using a micromanipulator (for coordinates and volumes see below). The needle was left in place for 10 min after injection. Viral injections were unilateral for optogenetic, fiber photometry, slice-physiology connectivity and rabies mapping experiments. For DREADD and taCasp3 experiments the viral injections were bilateral. For optogenetic and fiber photometry experiments, mice were also implanted with a unilateral fiber optic cannula secured to the skull with dental cement. Fiber optic cannulas were 200 μm for optogenetic and fiber photometry experiments. We used the following stereotactic coordinates (in mm): NAc (AP +1.35, ML ±1.35, DV −5.05 (virus) and –4.85 (fiber optic)), aBNST (AP +0.20, ML +0.80, DV –4.05 (virus)).

Adeno-associated viruses (AAVs) carrying Cre-inducible (double-inverse orientation; DIO) transgenes were packaged in our laboratory (AAVs for optogenetics, DREADD, taCasp3) or purchased from BrainVTA (Wuhan, China. http://brainvta.com) (AAVs for retrograde tracing or fiber photometry). Glycoproteindeleted rabies virus for retrograde tracing (RV-ENVA-ΔG-dsRed 2.0× 10^8 IFU/mL) was also purchased from BrainVTA.

### *In vivo* anesthetized electrophysiology

Adult mice (8-12 weeks) were anesthetized with isoflurane (4.0% for induction and set-up on the animal bed, 0.8%-1% during experiments) in a 20% O_2_/80% air mixture). Once anesthetized, mice were placed into a stereotactic frame and body temperature was maintained at ∼ 37 °C using a heating pad. A recording electrode was implanted into the NAc shell and a reference electrode was implanted in the contralateral NAc shell. Optical stimulation-induced neuronal activity was measured by calculating the firing rate 10 s before the stimulus and 40 s during the stimulus and the time bin of the firing rate was set to 500 ms. To determine when a single unit significantly responded to the optical stimulus, we used the criterion that the unit’s *P*-value of the ranksum test needed to be less than 0.05. The average firing rate of each group of neurons was calculated after normalizing the firing rate of each unit by Z-score method. To further examine the optical response for the classified neurons, the optical-induced firing probability were also calculated with the parameters of 15 ms pre-time, 15 ms post-time, and 0.1 ms time bin.

### Patch-clamp electrophysiology

Coronal slices (300 μM) containing NAc shell (bregma 1.7 to 0.6 mm) were prepared from *PV-Cre* transgenic mice using standard procedures. Brains were quickly removed and chilled in ice-cold modified artificial cerebrospinal fluid (ACSF) containing (in mM): 110 Choline Chloride, 2.5 KCl, 1.3 NaH_2_PO_4_, 25 NaHCO_3_, 1.3 Na-Ascorbate, 0.6 Na-Pyruvate, 10 Glucose, 2 CaCl_2_, 1.3 MgCl_2_. NAc slices were then cut in ice-cold modified ACSF using a Leica vibroslicer (VT-1200S). Slices were allowed to recover for 30 min in a storage chamber containing regular ACSF at 32∼34 °C (in mM): 125 NaCl, 2.5 KCl, 1.3 NaH_2_PO_4_, 25NaHCO_3_, 1.3 Na-Ascorbate, 0.6 Na-Pyruvate, 10 Glucose, 2 CaCl_2_, 1.3 MgCl_2_ (pH 7.3∼7.4 when saturated with 95% O_2_/5% CO_2_), and thereafter kept at ∼25 °C until placed in the recording chamber. The osmolarity of all the solutions was 300∼320 mOsm/Kg.

For all electrophysiological experiments, slices were viewed using infrared optics under an upright microscope (Eclipse FN1, Nikon Instruments) with a 40x water-immersion objective. The recording chamber was continuously perfused with oxygenated ACSF (2 ml/min) at 34 °C. Pipettes were pulled by a micropipette puller (Sutter P-2000 Micropipette Puller) with a resistance of 3-5 MΩ. Recordings were made with electrodes filled with intracellular solution (in mM): 130 potassium gluconate, 1 EGTA, 10 NaCl, 10 HEPES, 2 MgCl_2_, 0.133 CaCl_2_, 3.5 Mg-ATP, 1 Na-GTP. Inhibitory postsynaptic potentials (IPSPs) were recorded with PV cells held at 30 pA and the recording electrodes (7–9 MΩ) were filled with a solution containing (in mM): 120 cesium methansulphonate, 20 HEPES, 0.4 EGTA, 2.8 NaCl, 5 tetraethylammonium chloride, 2.5 MgATP, 0.25 NaGTP (pH 7.4, 285 mOsm/kg). AP-5 (25 μM) and NBQX (25 μM) were added to the ACSF during IPSP recording. IPSPs were blocked by adding 20 μM bicuculline (GABA_A_ receptor antagonist). Action potential firing frequency was analyzed in current-clamp mode in response to a 2 s depolarizing current step. Rheobase was determined as the amplitude of a minimum current step (advanced in 10 pA increments) to elicit an action potential response. All recordings were conducted with a MultiClamp700B amplifier (Molecular Devices). Currents were low-pass filtered at 2 kHz and digitized at 20 kHz using an Axon Digidata 1440A data acquisition system and pClamp 10 software (both from Molecular Devices). Series resistance (Rs) was 10∼30 MΩ and regularly monitored throughout the recordings. Data were discarded if Rs changed by >25% over the course of data acquisition.

### Measurements of norepherine (NE) and Corticotropin-releasing hormone (CRH)

Blood samples were collected from naive and stressed mice. Serum was prepared after each blood sample was centrifuged at 1000 rpm. Serum aliquots were immediately frozen at −80 °C prior to being used. NE and CRH in the serum were determined using the Radioimmunoassay Kits (Xinfan Biotechnology Co., Ltd, Shanghai, China) in accordance with the manufacturer’s instructions.

### Histology and confocal microscopy

Mice were deeply anesthetized and transcardially perfused with ice-cold 4% paraformaldehyde (PFA) in PBS (pH 7.4). Brains were fixed overnight in 4% PFA solution and then equilibrated in 30% sucrose in PBS. Next, 30 μm coronal slices were cut using a freezing microtome. Slices were stored in a cryoprotection solution at 4 °C until further processed. The sections were incubated with primary antibodies overnight at 4 °C. Alexa Fluor^®^488, 594 or 647-conjugated goat anti-rabbit or anti-mouse IgG antibodies (1:500; Invitrogen, CA, USA) were the secondary antibodies. Nuclei were counterstained using DAPI. Immunostaining was performed using mouse anti-parvalbumin (1:300, Millipore, MA, USA), rabbit anti-choline acetyltranserase (1:200, Abcam, ab6168). Histological slides were imaged on a (Zeiss LSM880) confocal microscope using a 10x, 20x or 63x objective.

### Retrogradely monosynaptic tracing

For monosynaptic retrograde tracing, 100 nl of mixed helper AAV (Ef1a-DIO-His-EGFP-2a-TVA-WPRE-pA and Ef1α-DIO-RVG-WPRE-pA) (BrainVTA) was injected into the NAc shell of PV-Cre transgenic mice using coordinates of +1.35 AP, +1.35 ML, and -5.05 DV. Three weeks later, rabies virus EnvA-pseudotyped RV-ΔG-DsRed (200 nl) (BrainVTA) was injected into the NAc shell using the same coordinates. Approximately seven days after the second injection, mice were anaesthetized with 10% chloral hydrate (0.4 mg/kg) and transcardially perfused with PBS, then 4% PFA (wt/vol) and then brain slices were prepared for tracing with dsRed.

### *In situ* hybridization

We used single-probe in situ hybridization on fixed frozen sections. Coding region fragments of Gad1, Gad2, Vglut1, Vglut2 and CRH were isolated from mouse brain cDNA using PCR and cloned into the pCR4 Topo vector (Thermo Fisher). Digoxigenin (DIG)-labeled riboprobes were prepared using a DIG RNA Labeling Kit (11277073910, Roche). Brain sections were hybridized to DIG-labeled cRNA probes at 56 °C for 14-16 hr. After hybridization, sections were washed twice in 0.2 x SSC at 65 °C for 20 min and then incubated with horseradish peroxidase (POD)-conjugated sheep anti-DIG antibodies (1:300; 1207733910, Roche) diluted in blocking buffer (1% Blocking reagent, FP1012, Perkin Elmer) for 45 min at room temperature (RT). Sections were washed three times for five minutes at RT in PBST (0.05% Tween 20 in 1 X PBS) wash buffer, and then treated using a TSA-plus Cy5 kit (1:100; NEL745001KT, Perkin Elmer) for 10 min at RT. Sections were washed two times for five minutes at RT in PBST and then incubated with Anti-RFP antibody (1:200; ab62341, Abcam) for 1.5 hr at RT, and washed. Sections were incubated with Alexa Fluor 594-conjugated goat anti-rabbit IgG antibodies (1:200; 115-587-003, Jackson Immuno Research) for 1 hr at RT. Sections were mounted in Fluoromount-G (0100-20, Southern Biotech) and then imaged using LSM 880 confocal microscopes (Zeiss, LSM880).

## STATISTICAL ANALYSIS

All statistical parameters for specific analyses are described in the appropriate figure legends.

### EPM score

ANOVAs were used to identify neurons regulated by the EPM arm types and were calculated using the firing rate of each neuron with a three level factor (closed arms, open arms and center zone) after each neuron’s spike train was binned into 3 s bins^53^. A neuron’s firing rate was considered to be influenced by EPM position if the rate in one maze area was statistically significantly higher than that in the others (closed arms versus open arms and center zone, open arms versus closed arms and center zone, center zone versus closed arms and open arms, Bonferroni post hoc test, \*\*\**P* < 0.001)^53^. EPM scores were used to quantify the degree to which a neuron can distinguish the structure of the EPM^49, 53^; EPM scores were calculated as previously described^49, 54^.

Score = (A-B) / (A+B), where

A = 0.25 × (|F_L_-F_U_| + |F_L_- F_D_| + |F_R_ - F_U_| + |F_R_ - F_D_|) and

B = 0.5 × (|F_L_ - F_R_| + |F_U_-F_D_|).

Horizontal and vertical arms represent closed and open arms, respectively. F_R_, F_L_, F_D_ and F_U_ are the % differences from mean firing rate in right, left, down and up arms, respectively; A is the mean difference in normalized firing rate between different-type arms and B is the mean difference for same-type arms. Neurons with firing patterns related to the EPM task have a high EPM score, as neurons will have similar firing rates in the same arm types (resulting in a small B value) and large differences in rates between different arm types (resulting in a large A value). The maximum EPM score of 1.0 shows no difference in firing rate across arms of the same type (B = 0). Negative EPM scores show that firing rates were more similar across arms of different types than across arms of the same type.

We calculated whether there was a statistically significant difference between the population of experimentally observed EPM scores from that expected by chance using a bootstrapping method. For each unit that had n spikes, 500 simulated EPM scores were generated by calculating the EPM score of n randomly chosen time stamps 500 times. 500 × 98 EPM scores were generated from 98 units recorded. Statistical differences between experimentally observed EPM scores of all neurons and chance were calculated by comparison to the simulated distribution of EPM scores using the Wilcoxon rank-sum test^49, 53^.

The firing pattern of NAc neurons at transitions between different types of EPM arms and Z-scores of firing rate were calculated for each unit for 10 s and averaged over total transitions for each unit. We identified a point where there was a change in the slope of the averaged z-scores. The averaged z-scores were divided into two parts by using this identified change point and a nonparametric Kolmogorov-Smirnov test was used to evaluate whether there were statistically significance differences between the means from these two data segments. This was calculated using 0.25 s bins.

### *In vivo* calcium signal analysis

Within each heat map, every row was normalized from 0 to 1 according to the formula (*D*-*D*_min_)/ (*D*_max_-*D*_min_), where *D* is the raw fiber photometry signal data, *D*_min_ is the minimum value of a given row and *D*_max_ is the maximum value of the same row. We sorted every row according to the time of peak value from late to early. To compact the heat map, we inserted one thousand data points between every two points of raw data by applying cubic spline methods. The raw heatmap data were normalized by Z-Score normalization. The formula for Z-Score is (*D*-μ)/ σ, where *D* is the raw fiber photometry signal data, μ is the mean value of raw data and σ is the standard deviation of raw data. The Z-Score data was divided into two even parts by time zero (defined by the time that mouse moved from closed arm to center zone). To visualize the mean recording traces from the activity of different NAc neuronal populations in the EPM, the first three seconds was chosen as a baseline and then Z-Score normalization was applied to all data using theΔ*F*/*F =* (*F-F_0_*)*/F_0_* method, where *F* is the normalized fiber photometry signal data and *F_0_* is the mean normalized data value. All calculations were performed using MATLAB 2017a, GraphPad Prism7 and SPSS 18.

### Single-unit spike sorting and analysis

Single-unit spike sorting was performed using Offline sorter (Plexon). Wideband signals were high-pass filtered (300 Hz) with a Bessel filter for detection of the spikes. The threshold value for spike detection was -4.5 standard deviations and spike waveforms were recorded for a time window of 1400 μs starting 300 μs before threshold crossing. Principal component values were calculated for the unsorted aligned waveforms and plotted in three-dimensional principal-component space. A group of waveforms were considered to be generated from one single unit if the waveforms were distinct from other clusters in the principal-component space and exhibited a refractory period more than 1 ms. In order to avoid analysis of the same neuron on different channels, cross-correlation histograms were calculated: if a neuron showed a peak at the same time as the reference neuron fired, only one of the two neurons was reserved for further analysis. To quantify the separation between identified neurons, L ratio and Isolation Distance ^55^ were calculated. High values of the Isolation Distance and low values of the L ratio indicated good cluster separation. The L ratio estimates the degree of noise contamination of one cluster, and a smaller value implied a lower degree of contamination. The Isolation Distance measured the average distance expected between a cluster and an equal ensemble of spikes outside the cluster, and a bigger value indicated a well-isolated cluster. The threshold values of the L ratio and Isolation Distance were set to 0.2 and 15, respectively^56^. Units with L ratio higher than 0.2 and Isolation Distance lower than 15 were excluded from the following analysis. Classification of NAc neurons were as described in previous studies and two features used for this, peak-to-peak width and average firing rate, were calculated for each unit^29, 57^. An unsupervised cluster algorithm based on Ward’s method was used to classify the neurons. Euclidian distance was calculated between neuron pairs based on the two-dimensional space defined by the two features^58^. To calculate a neuron’s burst number, a burst was defined as comprising at least three spikes with interspike intervals < 9 ms^59^. Neuron firing rates were considered as having undergone a statistically significant change if the P-value of the *ranksum* test was less than 0.05. The multitaper method^60^ in the Chrounux analysis package (http://chronux.org) was used for power spectra, time frequency and coherence analysis. The value was calculated using a 1 s window, 3 time-bandwidth product (NW) and 5 tapers. The coherence value in the theta band (5∼9 Hz) that exceeded the 95% confidence level was used for the analysis. The coherence value was normalized by dividing by the maximum value in the theta band. The statistical analysis of the coherence was conducted on the original values.

## Results

### Chronic stressors increase functional brain connectivity between the BNST and NAc

To gain a circuit-level understanding of anxiety-related behavior, we adopted a chronic stress model to investigate specific brain regions in the regulation of avoidance behavior under anxious states.

After exposure to chronic stressors (Fig. 1a), mice showed higher anxious states by markedly avoiding the center in open-field test (OFT) and open-arms of an elevated plus maze (EPM) when compared to their naive littermates (Fig. 1b-c). Stress related hormones, corticotropin releasing hormone (CRH) and norepinephrine (NE), were both significantly higher in chronically stressed mice (Fig. 1d). These data indicate that chronic stressors disrupted normal anxiety-related behavior and resulted in a maladaptive and excessive avoidance coping response.

**Fig 1.**
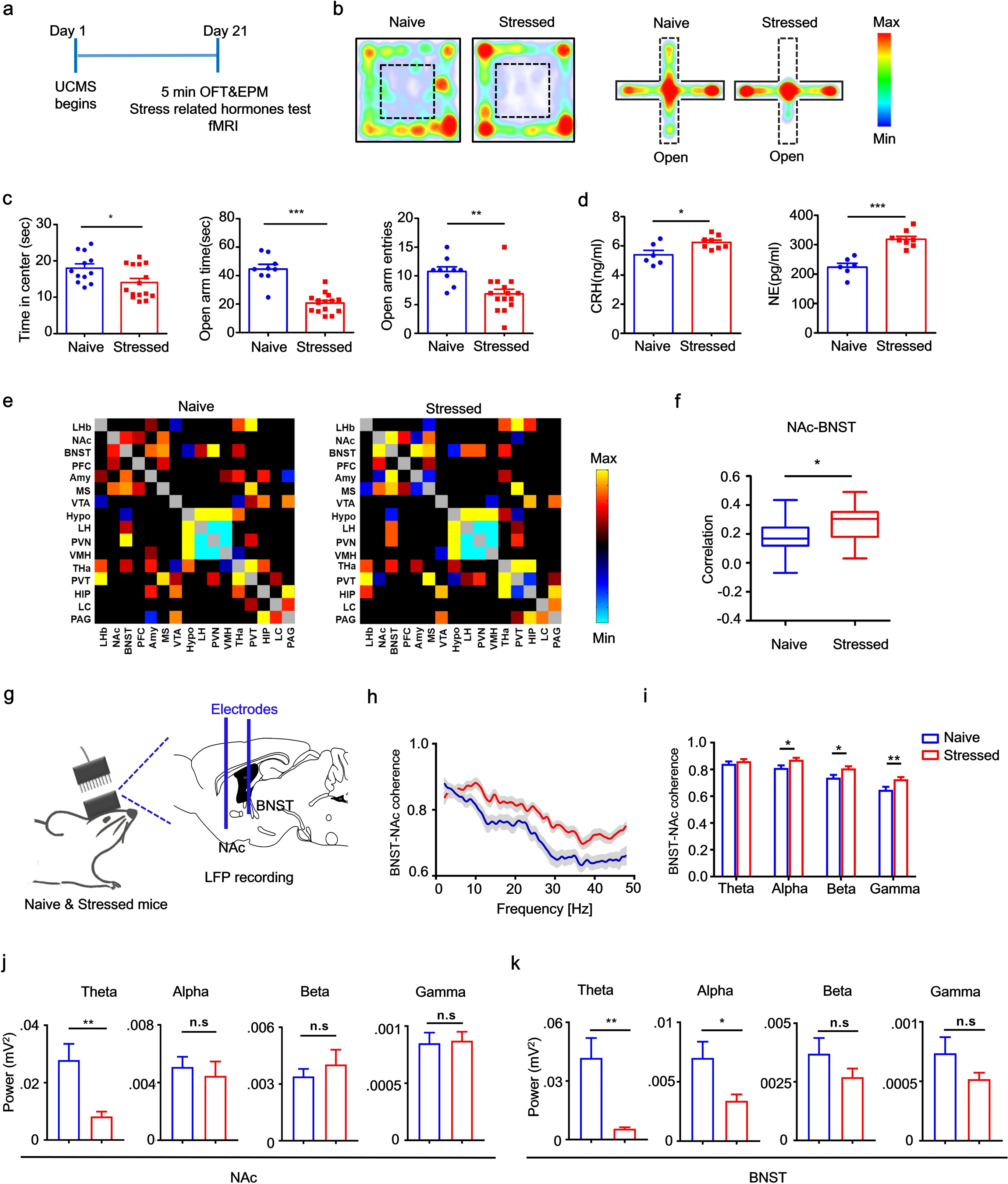
Chronically stressed mice exhibited increased functional brain connectivity in the BNST and NAc. (a) Protocol used for unpredictable chronic mild stress (UCMS) and functional brain connectivity measurement and paradigm for behavioral assay. (b) Representative mouse trajectory map for both naive and stressed groups in the open field test (OFT) and elevated plus maze (EPM); warm colors represent high time spent. (c) *Left*, time spent in central compartment; *middle*, time spent in the open arms; *right*, open-arm entries of naive control (n=9) and stressed groups (n=15) (Unpaired *t* test, *left*, *t* = 2.401, *P* = 0.0241; *middle*, *t* = 7.145, *P* < 0.0001; *right*, *t* = 3.214, *P* = 0.0040). (d) *Left* serum CRH and *right* NE concentrations of naive control (n=6) and stressed groups (n=8) (Unpaired *t* test, *left*, *t* = 2.671, *P* = 0.0204; *right*, *t* = 5.865, *P* < 0.0001). (e) Correlation matrices derived from global fMRI BOLD signal analysis (pseudocolor map of *t* statistics after thresholding at a false discovery rate, q of 0.05) across brain regions in stressed compared to wild-type naive littermates; warm colors represent higher correlations. (f) Correlation of BOLD synchronization in BNST-NAc in stressed (n=7) and naive littermates (n=5) (Unpaired *t* test, *t* = 2.698, *P* = 0.0108). (g) Schematic showing local field potential recording strategy. (h) Coherence between LFP signals recorded from NAc and BNST in stressed (n=7) and naive control mice (n=7). (i) Coherence in *theta* (4-8 Hz), *alpha* (8-12 Hz), *beta* (12-30 Hz) and *low gamma* (30-50 Hz) bands in NAc and BNST (Unpaired *t* test, *Theta*: *t* = 0.8605, *P* = 0.3934, *alpha*: *t* = 2.512, *P* = 0.0151, *beta*: *t* = 2.584, *P* = 0.0126, *low gamma*: *t* = 2.856, *P* = 0.0061). (j-k) Sum of power spectra obtained for LFP recordings in NAc shell (Unpaired *t* test, *theta*: *t* = 2.974, *P* = 0.0058, *alpha*: *t* = 0.5235, *P* = 0.6045, *beta*: *t* = 0.764, *P* = 0.4509, *low gamma*: *t* = 0.1712, *P* = 0.8652) and BNST (Unpaired *t* test, *theta*: *t* = 3.114, *P* = 0.004, *alpha*: *t* = 2.226, *P* = 0.0337, *beta*: *t* = 1.203, *P* = 0.2383, *low gamma*: *t* = 1.341, *P* =0.1901). \**P* < 0.05, \*\**P* < 0.01 and \*\*\**P* < 0.0001. Error bars (c-d, f, i, j, k) are mean±SEM. In H, the curves and shaded areas indicate the mean±SEM.

We further tested global functional brain connectivity by quantifying the synchronization of blood oxygen level-dependent (BOLD) fMRI signals across brain regions in anesthetized naive and chronically stressed mice. The synchronization of BOLD signals in the BNST and NAc was significantly higher in chronically stressed mice compared to their naive littermates (Fig. 1e-f). Moreover, functional connectivity was higher in stressed mice compared to their naive littermates between the basolateral amygdala (BLA) and BNST, and lower between the periaqueductal gray (PAG) and locus coeruleus (LC) (Fig. S1a, left and middle), whereas connectivity between NAc and PFC was not significantly different between groups (Fig. S1a, right). Consistent with the fMRI synchronization data, fMRI heat maps in coronal sections generated from stressed mice show a higher correlation of resting-state fMRI BOLD signal than those generated from their naive littermates, with a seed in NAc across brain regions (Fig. S1b, top) between BNST and NAc (Fig. S1b, bottom). To quantify long-range functional connectivity, we measured local field potential (LFP) coherence between the BNST and NAc in awake behaving stressed and naive mice (Fig. 1g). Compared to naive littermates, the stressed mice showed higher BNST-NAc coherence in low *gamma*, *alpha* and *beta* bands, but did not show any significant coherence in *theta* band in these two structures (Fig. 1h-i). We then analyzed the local *alpha*, *beta*, *gamma* and *theta* rhythms, respectively, in these two brain regions and found that local *theta* power was significantly lower in stressed mice compared to their naive littermates in both structures, whilst there was a trend towards lower *gamma* power in stressed mice in the BNST (Fig. 1j-k).

Since in the cortex and hippocampus, changes either in *theta*^25, 26^ or in *gamma* power^27^ is due to the PV cell functions, we then investigated PV neuronal firing features within the BNST-NAc circuit in chronically stressed mice.

### Chronically stressed mice have higher sNAc^PV^-neuron firing rates

Next, we looked at the distribution of PV neurons in both the BNST and NAc and found that PV neurons were expressed predominantly in the NAc shell (sNAc, Fig. 2a). Consistent with other work^28^, we found no expression of PV soma in the BNST, only terminal structures (Fig. 2b). To further investigate the causal relationship of PV neuronal activity and excessive avoidance observed in stressed mice, we tested sNAc^PV^ neuronal firing properties in stressed mice. We selectively expressed ChR2-mCherry in sNAc^PV^ cells of PV-Cre mice to visualize the PV neurons and recorded their firing patterns in response to electrical stimulation (Fig. 2c). PV neurons in stressed mice exhibited an increase in firing frequency in response to injection currents (Fig. 2d-f) without changes in resting membrane potential (RMP) or threshold (Fig. 2g-h). If excessive avoidance in stressed mice is due to increased excitability of the sNAc^PV^ neurons, it may be possible to rescue this maladaptive behavior by inhibiting sNAc^PV^ neuronal activity. To test this possibility, we injected AAV-PV-Cre and either DIO-hM_4_D_i_ or DIO-mCherry into the sNAc of stressed mice (Fig. 2i). Compared to the mCherry group, chemogenetic inhibition of PV neurons rescued the excessive avoidance effect observed on the open arms of EPM, as seen in both open arm time and entries, which were significantly higher (Fig. 2j-k). These results imply that hyper-excited sNAc^PV^ neurons in stressed mice contributed to their excessive avoidance behavior.

**Fig 2.**
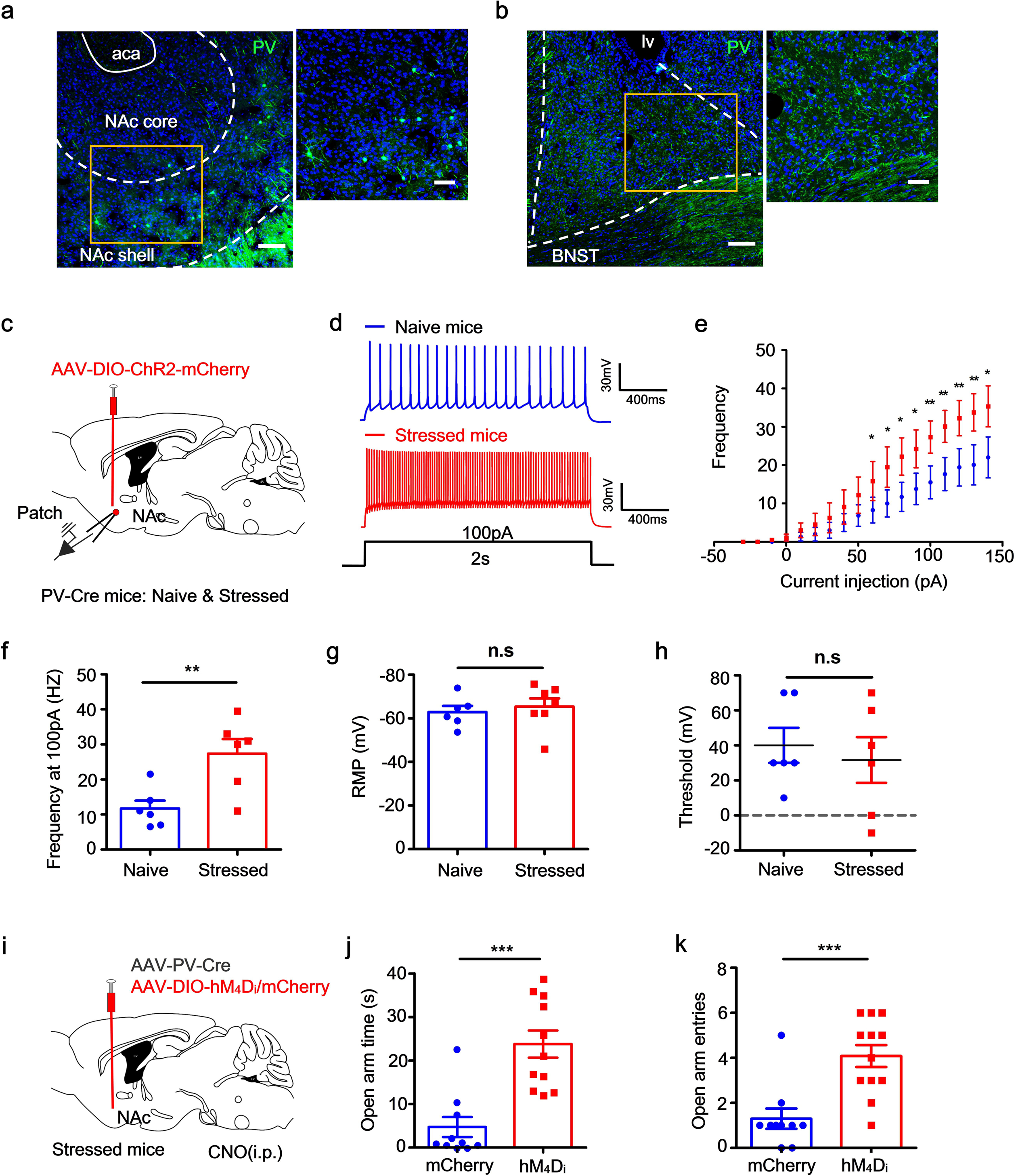
Accumbal PV neurons exhibit higher excitability in chronically stressed mice, and are responsible for excessive avoidance of anxiogenic locations. (a-b) Representative images showing the distribution of PV neurons (green) in NAc and BNST; *left*, scale bar, 100 μm; *right*, scale bar, 50 μm. (c) Schematic showing selective expression of ChR2 into the sNAc^PV^ neurons for whole-cell patch clamp recordings. (d) Representative traces showing action potential spiking in sNAc^PV^ neurons (n=4 mice per group). (e) The input–output curve of injected currents versus spiking frequency. (f) Spiking frequency of PV neurons in naive and stressed mice during the injection of a current (100 pA, n = 6-8 cells from 4 mice per group, unpaired *t* test, \**P* < 0.05; \*\**P* < 0.01). (g-h). RMP and threshold of naive and stressed mice (n = 6-8 cells from 4 mice per group, unpaired *t* test, \**P* < 0.05; \*\**P* < 0.01). (i) Schematic showing chemogenetic inhibition of sNAc^PV^ neurons. (j-k) Mean time spent in open arms and entries to the open arms with or without chemogenetic inhibition of sNAc^PV^ neurons (n = 10-12 mice per group, unpaired *t* test, \**P* < 0.05; \*\**P* < 0.01; \*\*\**P* < 0.001).

### PV neurons in sNAc represent an anxiety-related signal and impact avoidance coping behavior

We next investigated how NAc neurons are engaged in anxiogenic information processing under physiological conditions using a combination of single-unit and photometry recordings in freely-moving mice. These approaches allowed us to record neuronal firing events in the NAc shell whilst mice freely explored safe/threatening environments. Ninety-eight well-isolated NAc units from nine mice during the EPM assay were recorded. Distinct sub-types of NAc neurons were classified based on their major electrophysiological properties^29, 30^. Neurons were classified as: 1) putative fast spiking units (FS) if mean firing rate was more than 15 Hz, the initial slope of valley decay was greater than 22 mv/μs and the valley half decay time was less than 250 μs; 2) putative non-fast spiking units (Non-FS) if the firing rate was less than 2 Hz, the initial slope of valley decay was less than 22 mv/μs, and the valley half decay time was greater than 250 μs; 3) others (units that could not be classified as FS or non-FS) or 4) unclassified, the units could not be identified as neurons (Fig. 3A). A total of five putative FS neurons (5.1%) and 28 putative Non-FS neurons (28.6%) were clearly identified (Fig. 3a).

**Fig 3.**
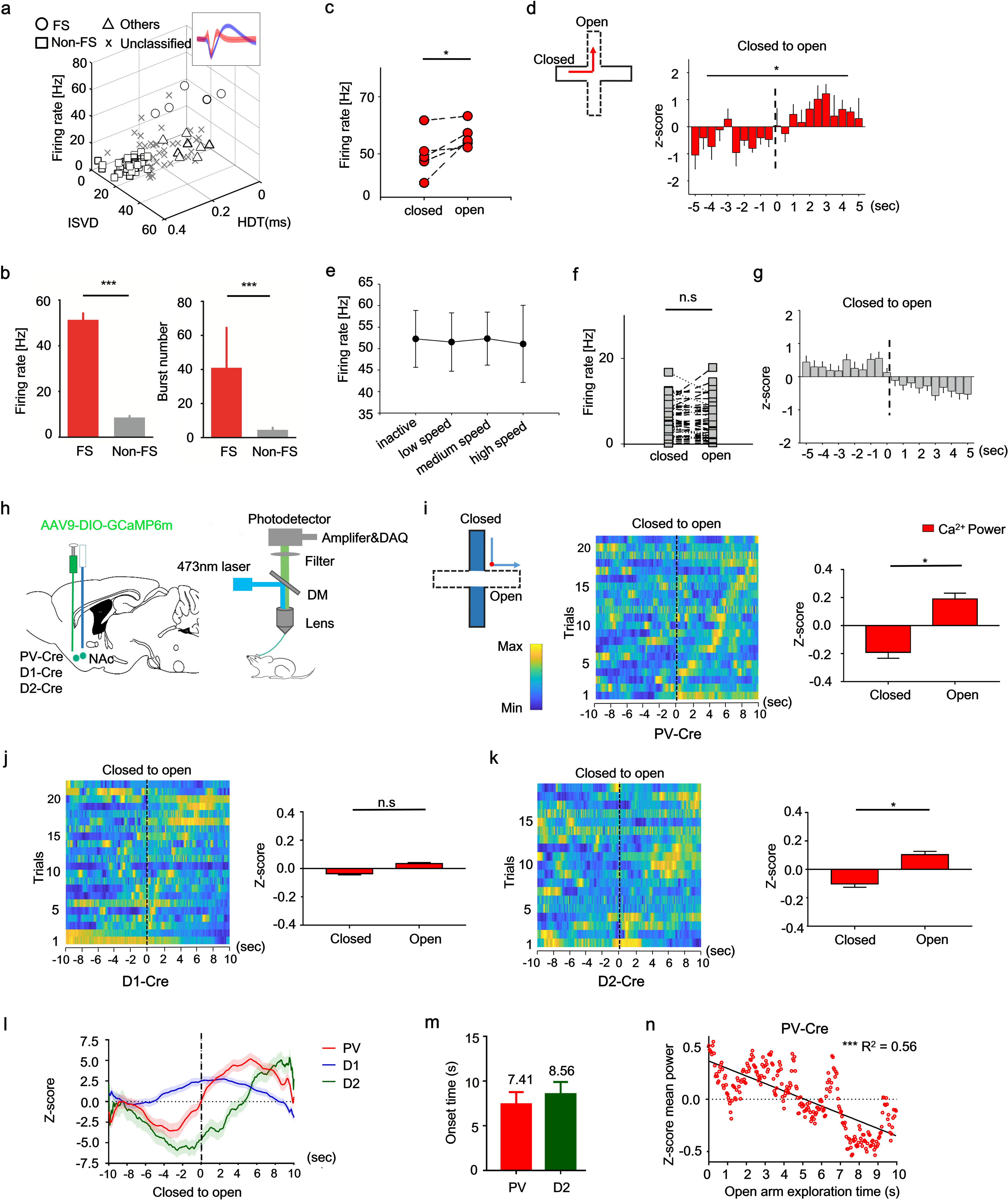
Different neuronal populations in the NAc shell show distinct firing preference before and during exploration of anxiogentic environments. (a) Scatter plot of the firing rate and peak-to-peak width for 98 units from 9 mice. *Inset*, representative waveforms from an identified Non-FS (blue) and FS (red) neuron; ISVD, initial slope of valley decay; HDT, valley half decay time. (b) *Left*, mean firing rates of FS and Non-FS neurons (Mann-Whitney rank sum test, n of FS = 5, n of Non-FS = 28, *t* = 155, \*\*\**P* <= 0.001); *right*, burst number of FS and Non-FS neurons (Mann-Whitney rank sum test, n = 5 (FS), n = 28 (Non-FS), *t* = 155, \*\*\**P* <= 0.001). (c) Scatter plot showing the firing rate of FS neurons between closed and open arms (Paired *t* test, n = 5, *t* = -2.620 with 4 degrees of freedom, *one-tailed *P* = 0.032). (d) Z-scores of FS neurons during movement from the closed to open arms (nonparametric Kolmogorov-Smirnov test, n = 5, \**P* < 0.05, see in Methods); *inset,* horizontal and vertical arms represent closed and open arms, respectively. (e) Correlation between locomotion and firing rates of the FS neurons. (f) Scatter plot showing the firing rates of Non-FS neurons in closed and open EPM arms (Paired *t* test, n = 28, *t* = 0.179 with 27 degrees of freedom, *P* = 0.86). (g) There was no difference in individual Non-FS neuron firing rates between closed and open arms. (h) Schematic showing fiber photometry recording strategy. (i-k) *Left*, normalized NAc Ca^2+^ activity maps from different neuron populations in the EPM (warm colors, high activity), binned by time (s) from the EPM crossing point (*inset*, red point); *inset,* horizontal and vertical arms represent open and closed arms, respectively; *right*, normalized NAc Ca^2+^ transients of different neuron populations in the EPM open arms compared to closed arms (Wilcoxon test, n = 250, \**P* < 0.05). (l) Mean activity traces from different NAc neuron populations during the EPM. (m) Comparison of the firing onset between PV and D2 cells when mice were approaching the crossing point in the EPM (n = 19 trials). (n) Normalized Ca^2+^ transients in NAc PV neurons was correlated with anxiety state (Open arm exploration time) (linear regression, F (1, 248) = 316.4, R^2^ = 0.56, *P* < 0.0001, n = 8 mice).

These two classes of neurons had significantly different mean firing rates and burst numbers (Fig. 3b, \*\*\**P* < 0.001, respectively). Individual FS units showed a firing preference for the open arms over the closed arms (Fig. 3c). The z-scores of FS unit firing rates increased when mice entered an open arm (Fig. 3d) and decreased when they moved to closed arms (Fig. S2a). Note that FS unit firing rates were not influenced by locomotion speed (Fig. 3e). By contrast, the individual Non-FS units showed no firing preference for either closed or open arms (Fig. 3f) and none of the z-scores of the firing rates were modulated by movement over the crossing point in the EPM (Fig. 3g, and see also Fig. S2b). We further checked the local *theta* power during exploration of either closed or open arms and found *theta* power was significantly lower during exploration of open arms compared to closed arms, implying an anxious state (Fig. S2c). There was no difference in theta power at different locomotion speeds (Fig. S2d). To show the temporal dynamics of the NAc *theta* oscillation on the EPM, a mean time-frequency map of mice traversing between arms was calculated and aligned to entrance time points. The map shows a decline in mean theta power when mice left the closed arms (Fig. S2e, *left*, indicated by the black dotted line), which increased when they returned to the closed arms (Fig. S2e, *right*, indicated by the black dotted line). Finally, to further address the relationship between local spike activity and neural oscillations, we calculated the mean spike-field coherence for both FS and Non-FS units. We found that FS, but not Non-FS, had strong coherence between their spikes and theta oscillations at 4-8 Hz (Fig. S2f, \*\**P* = 0.007). These results imply that NAc FS activity was inversely correlated with *theta* power, and that a reduction of the accumbal *theta* activity reflects a higher stress load during exploration of the threatening environments, which can promote adaptive avoidance behavior. These findings are consistent with the above result showing that a decrease in local *theta* power within either NAc or BNST is reflected by the anxious state of the stressed mouse (see also Fig. 1j-k).

PV neurons are fast spiking neurons^31–33^, and D1, D2-MSNs are the dominant non-fast spiking cell types in the NAc^34^. We next used *in vivo* calcium signal recordings in three different strains of transgenic mice to confirm the impact of these different neuron types on anxiogenic information processing (Fig. 3h). We first performed Immunostaining and *in situ* hybridization and confirmed the specific expression of GCaMP6_m_ in PV, D1R or D2R cells respectively (Fig. S3a-c). Ca^2+^ signals were then recorded as mice moved from the closed to open arms (Fig. 3i, inset). PV neurons within sNAc were activated when mice approached the boundary between the closed and open arms and Ca^2+^ signal increased significantly during exploration of the open arms (Fig. 3i); D1 MSNs exhibited no preference for either closed or open arms (Fig. 3j) whereas D2 MSNs showed a slightly different firing preference for the open arms to the closed ones (Fig. 3k). Recording traces from the different populations of NAc neurons were used to generate z-scores of calcium signal change and we found that PV and D2 neurons were active when mice were in the open arms but not the closed arms (Fig. 3l); however, the onset of D2 neuronal firing occurred later than the PV neurons during exploration of the open arms (Fig. 3m). This neural activity in the open arms may reflect that PV neurons in the NAc represent the association between anxiety-related information and the initiation of avoidance coping behavior. In addition, we found a well negative correlation between the PV firing rates and time of open-arm exploration (Fig. 3n), which suggests that accumbal PV neuronal activation plays a role in avoidance of anxiogenic locations. Calcium transients for from these cell populations in a negative control group are seen in Fig. S3d-f where no differences were found between open and closed arms.

### Activation of PV neurons in NAc shell is required for avoidance coping responses to anxiogenic stimuli

A combination of PV-Cre mice, conditional ChR2 viral expression, and optogenetic manipulation was used to test the impact of accumbal PV neuronal activity on avoidance behavior induced by an anxiogenic context. Because fast-firing PV cells can generate high-frequency trains with maximal frequencies greater than 100 Hz, and above 60-80 Hz the slope of frequency-interval (f-I) curve is well-approximated by linearization^35^, we used a photostimulation rate of 60 Hz for the optrode placed in the NAc shell (Fig. 4a, *left*); this is approximately the average firing rate of FS units in the open arms (see also Fig. 3c). Immunostaining results indicated that the majority of ChR2-mCherry labeled neurons expressed PV (Fig. 4a, *right*). We confirmed that opto-tagged PV^+^ cells, recorded from patch-clam experiments in the acute brain slices, were steadily activated by 5-80 Hz light stimulation at 470 nm (Fig. 4b). Selective light stimulation of these PV neurons in the NAc during the EPM task led to more avoidance behavior in PV-ChR2 mice compared to PV-mCherry controls, illustrated by a significantly lower number of entries and markedly less time spent in the anxiogenic open arms compared (Fig. 4c-d). The OFT led to similar results: light stimulation of PV neurons in PV-ChR2 mice led to less exploration time in the center of the open-field apparatus compared to the PV-mCherry control mice (Fig. 4e) without any difference in locomotion between the two groups during each five-min epoch (Fig. 4f).

**Fig 4.**
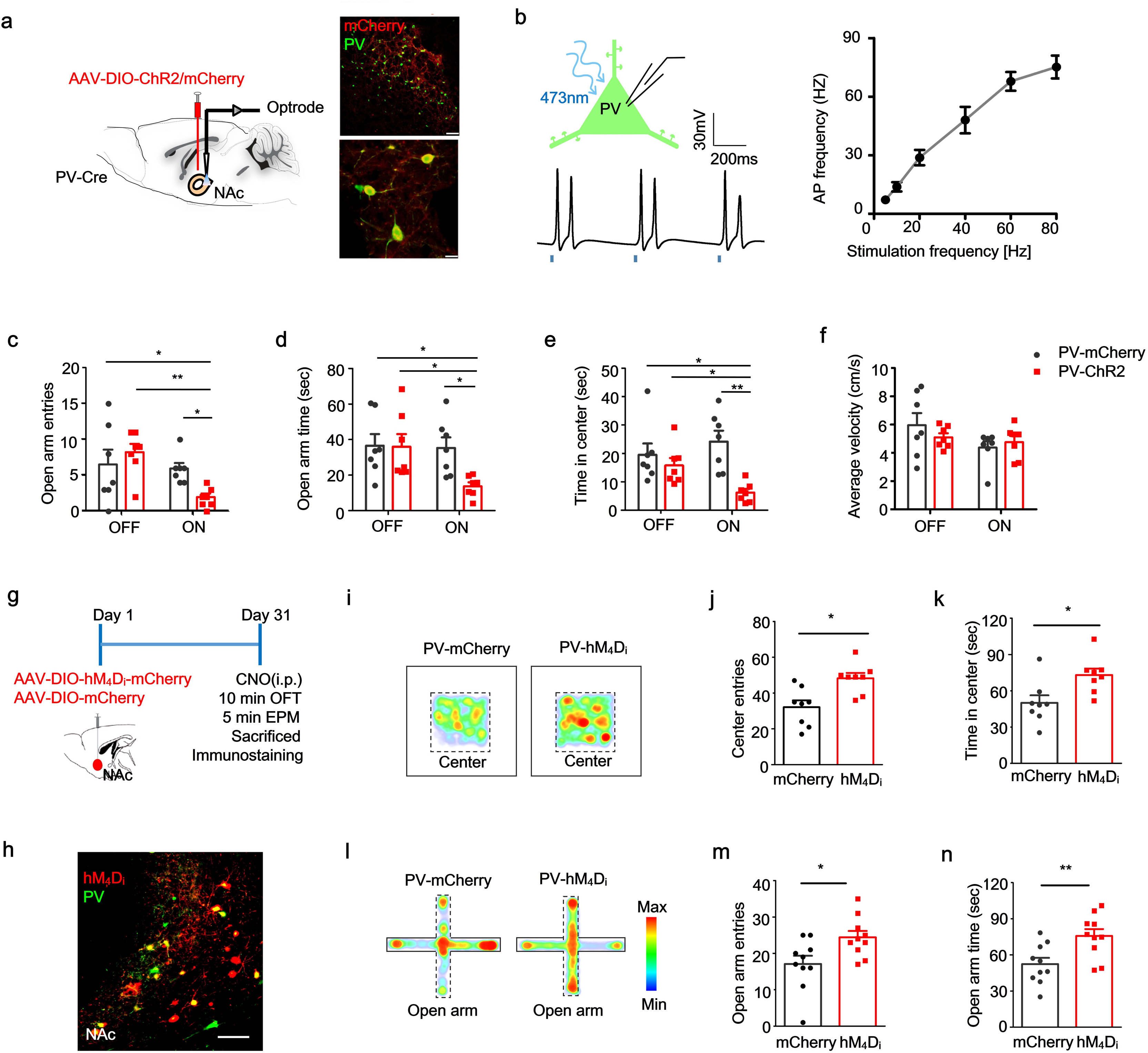
Activation of PV neurons in sNAc is required for anxiety-related avoidance. (a) *Left*, Schematic showing optrode placement in the NAc shell; *Top right*, ChR2-mCherry neurons (red) in NAc co-expressed PV (green); scale bar, 100 μm; *bottom right*, higher magnification of the co-staining (merged, yellow; scale bar, 10 μm). (b) *Top left*, schematic showing patch clamp technique; *bottom left*, sample of light-evoked PV neuronal action potentials; *right*, the action potential frequency of PV neurons following different light stimulation frequency from 5-80Hz. Comparison of (c) open arm entries and (d) time spent in the open arm between PV-ChR2 and PV-mCherry groups (n = 7, *t* test, \**P* < 0.05). (e) Comparison of time spent in the central area during OFT between PV-ChR2 and PV-mCherry groups (n = 7 per group, Two-way ANOVA, \*\**P* < 0.01, F3,18 = 5.99; Bonferroni post hoc analysis, \**P* < 0.05, \*\**P* < 0.01). (f) Mean velocity of PV-ChR2 and PV-mCherry groups during OFT (n = 7 per group, Two-way ANOVA, *P* = 0.1827, F (3, 18) = 1.8; Bonferroni post hoc analysis). (g) Protocol schematic for selective chemogenetic inhibition of PV neurons in the sNAc. (h) Immunostaining showing targeted hM_4_D_i_ expression (red) in PV neurons (green) of sNAc in PV-Cre mice; scale bar, 50 μm. (i) Representative mice thermal tracks for *left* PV-mCherry and *right* PV- hM_4_D_i_ groups during OFT. (j-k) Mean number of entries to center (n= 8, Paired *t* test, *t* = 2.829, df = 7, \**P* = 0.0254) and mean time in the center (n = 8, Paired *t* test, *t* = 2.448, df = 7, \**P* = 0.0442). (l) Representative mice thermal tracks for *left* PV-mCherry and *right* PV- hM_4_D_i_ groups during EPM task. (m-n) Mean number of open arm entries (n = 10-11, unpaired *t* test, *t* = 2.739, df =19, \**P* = 0.013) and mean time in open arms (n = 10-11, unpaired *t* test, *t* = 3.053, df = 19, \*\**P* = 0.0065).

In order to further determine the causal role of accumbal PV activity in avoidance behavior related to anxiety states, we expressed hM_4_D_i_ in PV neurons by bilateral injection of Cre-dependent AAV-DIO-hM_4_D_i_-mCherry into the NAc shell of PV-Cre mice. Mice were given a 10-min OFT followed by a 5-min EPM test (Fig. 4g). Co-localization of the majority PV cells with hM_4_D_i_ was verified by immunostaining (Fig. 4h). Representative OFT and EPM heat maps for both PV/mCherry (control) and PV/hM_4_D_i_ groups are shown in Fig. 4i-k: hM_4_D_i_ mice showed significantly greater center exploration, relative to mCherry controls. Consistent with the OFT findings, hM_4_D_i_ mice showed greater exploration of the open arms compared to the control mice, reflecting an inappropriate avoidance coping behavior (Fig. 4l-n).

Taken together, these data suggest that activity of PV neurons within the NAc is required for execution of appropriate avoidance behavior to buffer the stress response evoked by anxiogenic environments.

### Inputs to NAc^PV^ neurons originate predominantly from the anterior dorsal BNST (adBNST)

We next investigated whether the BNST was upstream of the NAc^PV^ neurons and if it is the region where anxiety-related avoidance coping behavior is integrated. Cre-dependent, rabies-virus-based whole brain monosynaptic tracing was performed to analyze upstream regions that innervate NAc^PV^ neurons. We injected PV-Cre mice with Cre-dependent AAVs expressing the avian EnvA receptor (TVA) and rabies virus envelope glycoprotein (RG) in combination with theΔG-dsRed (EnvA) rabies virus (RV) (Fig. 5a). We found that a Cre-dependent helper virus combined with RV expressing dsRed labeled 84% of NAc PV neurons (GFP^+^ cells, Fig. 5b). Our results showed that the dominant inputs to NAc^PV^ neurons were from the anterior dorsal part of the nucleus of the stria terminalis (adBNST) (52%, Fig. 5c, d), a classic GABAergic anxiety-associated region. Other brain regions that provided inputs to NAc^PV^ neurons included the basolateral amygdala (BLA, 4.34%), central amygdala (CeA, 10.27%), media prefrontal cortex (mPFC, 5.7%) as well as the ventral tegmental area (VTA, 6.07%). NAc^PV^ neurons also received monosynaptic inputs from reward-associated components, such as the paraventricular thalamus (PVT, 19.6%) and lateral hypothalamus (LH, 16.07%)^36^; no projections from the Hippocampus (Hippo, 0%) were found (Fig. 5c, d). *In situ* hybridization experiments demonstrated the co-expression of Gad 1/2 and dsRed in aBNST (96.1%, Fig. 5e, f). These findings indicate that NAc^PV^ neurons were modulated under a GABAergic network. No RV expressing dsRed labeled cells were found in the above brain regions (Fig. S4). We tested further and found no other neuronal markers in the BNST except sparse CRH co-expressed with RV signals in adBNST (Fig. S5).

**Fig. 5.**
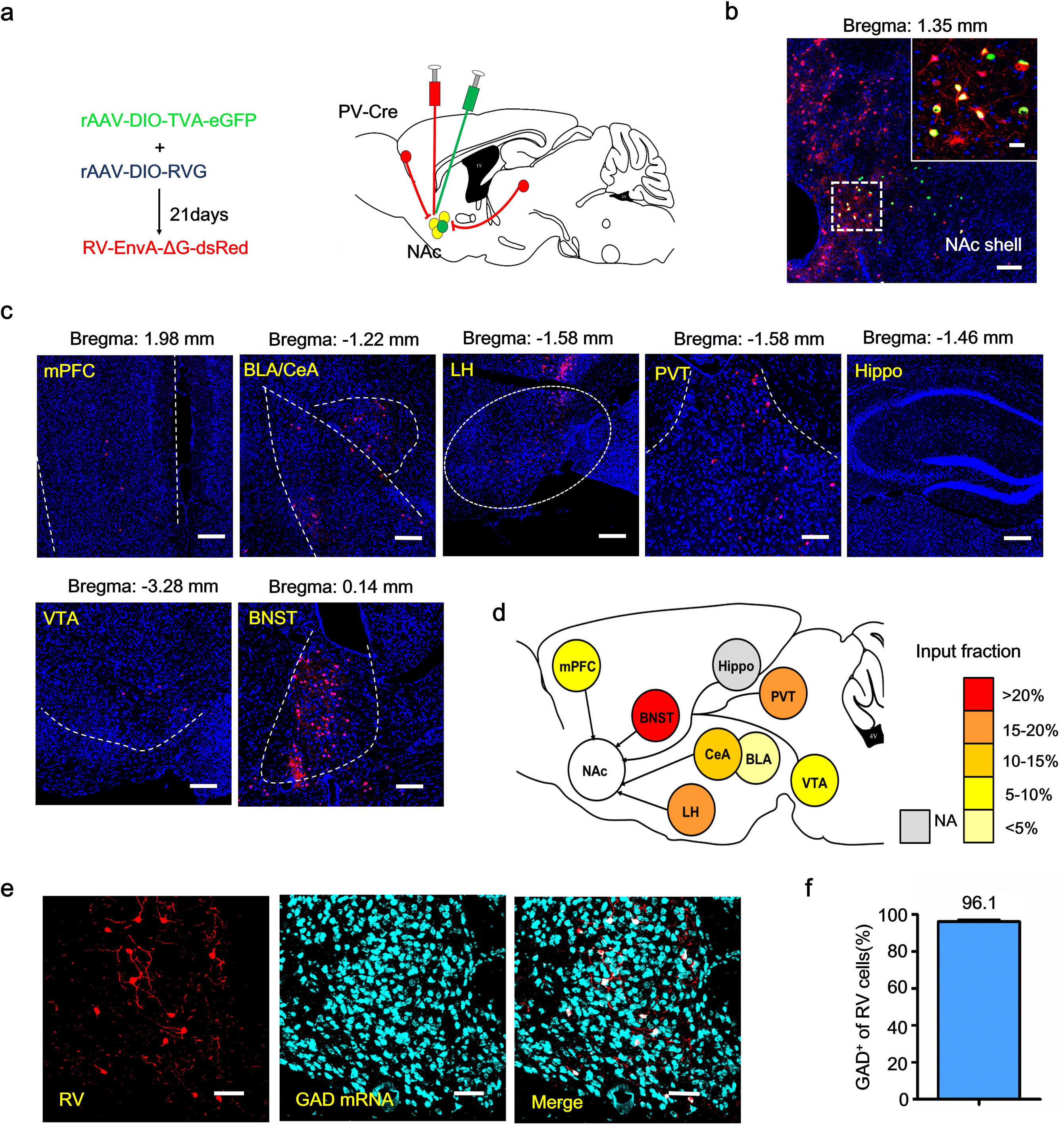
Monosynaptic GABAergic inputs to sNAc^PV^ neurons mainly stem from the adBNST. (a) Schematic showing sNAc injections of AAV-Ef1α-DIO-TVA-eGFP (AAV2/9) virus and AAV-Ef1α-DIO-RG (AAV2/9) on day 1 and RV-EvnA-DsRed on day 21 in PV-Cre mice to retrogradely trace input neurons (red) to NAc shell (yellow, starter neurons). (b) RV-mediated transsynaptic retrograde tracing of NAc inputs; fluorescence images of NAc region (coronal diagram) in PV-Cre mice (n = 3 mice); Scale bar, 100 μm; *inset*, enlarged view of the region in the dotted white box, showing starter cells (yellow, expressing both eGFP and DsRed, indicated by white arrowheads; scale bar, 50 μm.) (c) Typical coronal-section planes with distance (anterior-posterior) from bregma showing dsRed-expressing presynaptic neurons retrogradely labeled by NAc injection of virus in PV-Cre mice; scale bar, 100 μm; Hippo, hippocampus; BLA, basolateral amygdala; CeA, central amygdala; PVT, paraventricular thalamus; mPFC, medial prefrontal cortex; LH, lateral hypothalamus; VTA, ventral tegmental area; BNST, bed nucleus of the stria terminalis. (d) Overview of inputs to NAc PV neurons (n = 3 mice per region giving a total of 12-39 slices). (e) Sections were co-stained with antibodies against RFPn (RV) (red) and riboprobes (cyan) for *Gad1/2*; scale bar, 50 μm. (f) Approximately 96.1% of RV-infected neurons were co-labeled for *Gad1/2* in the adBNST (n = 12 slices, from 3 mice).

### Optogenetic activation of adBNST GABAergic terminals in sNAc produces an anxiolytic effect *via* NAc^PV^ neurons

Next, we investigated the impact of the connection between the adBNST GABAergic neurons and NAc^PV^ cells on producing anxiety-related behavior. We virally expressed GAD67-Cre and Cre-dependent channelrhodopsin-2 (ChR2) in adBNST GABAergic neurons and visualized NAc^PV^ neurons by injection of adeno-associated viruses (AAVs) encoding the fluorophore mCherry into PV-Cre mice (Fig. 6a). Co-staining results revealed that the majority of neurons expressing GAD^+^ also co-expressed ChR2 (Fig. 6b-c). We recorded evoked IPSPs from PV neurons within the NAc shell by illumination of adBNST afferent axon fibers, which was completely blocked by 20 μM bicuculline, implying a GABAergic monosynaptic input to the NAc PV neurons (Fig. 6d-e). The mean latency was 4.07± 0.7 ms, in line with monosynaptic transmission (Fig. 6f, right). These data suggest a direct functional GABAergic input to the sNAc^PV^ neurons. We further targeted the function of this GABAergic input to the NAc^PV^ neurons by virally expressing Cre-dependent ChR2 and GAD67-Cre in adBNST neurons followed by light stimulation of the terminals within sNAc (Fig. 6g). Blue light stimulation resulted in decreased avoidance of both EPM open arms and OFT center area (Fig. 6i). We then tested whether, under similar conditions, we would obtain similar results without adBNST neurons innervating NAc^PV^. AAV2/9-FLEX-taCasp3-TEVp and PV-Cre viruses were injected into the sNAc to selectively kill PV^+^ neurons; Cre-dependent ChR2 and GAD67-Cre were both injected into the adBNST of mice to specifically activate this new BNST-NAc GABAergic circuit (Fig. 6k); immunostaining confirmed that most of the PV^+^ neurons were killed by taCasp3 compared with the control virus (Fig. 6l). Terminals in the NAc shell were stimulated once again and we found that reduced avoidance of EPM open arms and OFT center was now effectively blocked during the light ON phase (Fig. 6m-n). These findings suggest a crucial role of adBNST GABAergic inputs to the NAc^PV^ neurons in distinguishing safety and risk to facilitate avoidance of anxiogenic locations. To examine the presynaptic effect of GABA release on the NAc^PV^ neurons, we recorded two consecutive eIPSPs, which were separated by varying interspike intervals to calculate the paired-pulse ratio (PPR), upon light stimulation of the adBNST GABAergic afferents to sNAc. We found that the PPR was significantly increased in stressed mice compared to naive ones at both 50- and 100-ms interstimulation interval (Fig. 6p-q). This increased PPR in stressed mice suggests an impaired presynaptic GABA release at adBNST to sNAc synapses, further indicating these sNAc PV neurons are disinhibited by GABAergic inputs from the adBNST under a chronic stress state.

**Fig. 6.**
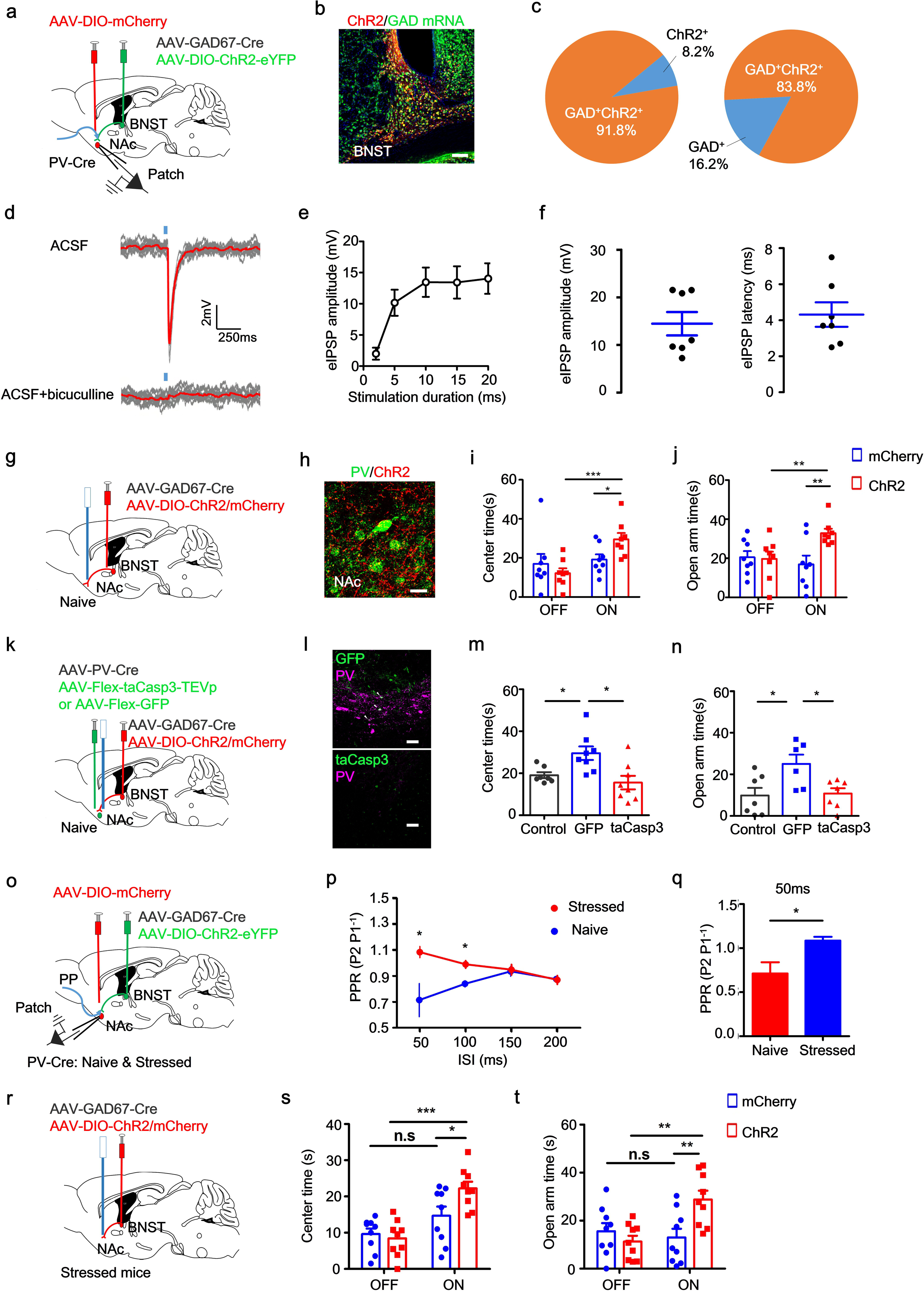
Optogenetic stimulation of adBNST GABAergic termimals in sNAc produces an anxiolytic effect, which is mediated by sNAc^PV^ neurons. (a) Schematic showing the strategy for selective optogenetic manipulation of GABAergic afferents from aBNST onto sNAc^PV^ neurons and recording postsynaptic inhibitory potentials (IPSP). (b-c) Representative image showing that most ChR2 positive cells in adBNST co-expressed GAD1/2 mRNA (n = 3 mice); scale bar, 100 μm. (d) *Top*, mean eIPSPs obtained from PV neurons within the NAc shell *(bottom*) was blocked by 20 μM bicuculline. (e) The eIPSP amplitude of PV neurons following different light stimulation duration from 1-20 ms. (f) Quantification of eIPSP amplitude (*left*, n = 7 cells from 4 mice) and latency (*right*, n = 7 cells from 4 mice). (g) Schematic showing optogenetic illumination of adBNST GABAergic afferent fibers that innervate NAc^PV^ neurons. (h) Representative image showing adBNST GABAergic afferent axons innervating the PV neurons within NAc shell. Scale bar, 10 μm. (i-j) Mean time spent in the center (n = 8 mice per group, unpaired *t* test, \**P* < 0.05; \*\**P* < 0.01; \*\*\**P* < 0001) and in open arms (n = 8 mice per group, unpaired *t* test, \**P* < 0.05; \*\**P* < 0.01; \*\*\**P* < 0001) with or without optogenetic stimulation of adBNST GABAergic afferent axons innervating PV neurons within NAc shell. (k) Schematic showing viruses injected into the adBNST and NAc shell. Three groups: Control, BNST^mCherry^-NAc^GFP^; GFP, BNST^ChR2^-NAc^GFP^; taCasp3, BNST^ChR2^-NAc^taCasp3^. In each mouse, an optical fiber was implanted above the NAc shell for optogenetic illumination of adBNST GABAergic outputs. (l) Representative images showing selective deletion of PV neurons in the NAc shell; Scale bar, 100 μm. (m-n) Mean time spent in the center (n = 6-7 mice per group, unpaired *t* test, \**P* < 0.05; \*\**P* < 0.01; \*\*\**P* < 0001) and in open arms (n = 6-7 mice per group, unpaired *t* test, \**P* < 0.05; \*\**P* < 0.01; \*\*\**P* < 0001). (o) Schematic showing PPR of eIPSPs recorded from PV neurons within the NAc shell in wild-type and stressed PV-Cre mice during optogenetic stimulation of adBNST GABAergic afferents around PV neurons. (p) PPR of eIPSPs plotted as a function of interspike intervals (ISIs); (q) mean PPR measured at 50-ms interspike interval. \**P* < 0.05 (n = 4 from 3 mice per group, *t* test). (r) Schematic showing optogenetic stimulation of adBNST GABAergic afferents around PV neurons in the sNAc on stressed mice. (s-t) Mean time in the center and open arms (n =9 mice per group, unpaired *t* test, \**P* < 0.05; \*\**P* < 0.01; \*\*\**P* < 0.001).

Based on these findings, we then tested whether activation of the adBNST GABAergic afferents to sNAc^PV^ could rescue the pathological anxiety-related behavior in stressed mice. We injected AAV-GAD67-Cre and either DIO-ChR2 or DIO-mCherry into the adBNST of stressed PV-Cre mice and implanted an optical fiber above the sNAc (Fig. 6r). Light stimulation of adBNST GABAergic afferents in sNAc significantly increased the time spent in the OFT center and open arms of the EPM (Fig. 6s-t), which implies that excessive avoidance behavior related to anxious state was rescued. Taken together, these findings indicate that in anxious states, the adBNST may send a disinhibition input to the sNAc^PV^, leading to excessive avoidance of the threatening locations.

## Discussion

Chronic stress leads to long-term changes in brain structure and function, which increases the incidence of stress-related disorders, such as anxiety^4^. Anxiety disorders are the most prevalent mental disorders and are associated with immense social health care costs^21^. A central symptom is avoidance behavior, which also acts as a reinforcer of the anxious state^24, 38^. It is vital then to understand the underlying cellular and circuitry mechanisms underpinning this type of avoidance behavior, which could result in new methods to break the cycle of anxiety-avoidance.

A critical role in stress response and anxiety has been attributed to the BNST in both rodent and human studies^3, 39, 40^. The NAc is a key component of the brain “reward” circuits^41–43^ in emotional and motivated actions and its dysfunction has been strongly implicated in emotional disorders^44, 45^, especially in exerting a dominant influence on anxiety ^10, 46, 47^. Although one study noted a projection from BNST to NAc^11^, knowledge about the function of the link between the stress response neurocircuitry and reward circuitry not known.

Our resting fMRI findings indicate that besides the high correlation among the stress response regions (Fig. S1a), increased connectivity was observed between BNST and NAc under a chronic-stress-induced anxious state (Fig. 1e-f); the LFP coherences in *alpha*, *beta* and *gamma* bands between these two regions were also significantly increased (Fig. 1i), confirming an intrinsic link between the stress response region and the reward circuit component under anxious state. We can envision that the increased synchronization of BOLD signals in BNST and NAc may potentially be used as an imaging marker for the diagnosis of anxiety disorders in the future.

LFP recordings further showed a marked decrease in the local *theta* power both in the NAc and BNST of stressed mice (Fig. 1j-k), indicating PV cell involvement within these two regions as previous studies have suggested a correlation between PV activity and local *theta* changes^25, 26^. However, only the NAc shell contained PV cell bodies (Fig. 2a) and the patch-clamp data found that these accumbal PV cells were hyper-excitable under an anxious state (Fig. 2d-e), confirming the surprising role of accumbal PV interneurons in behavior related to anxiety, given their low occurrence in the region. According to a previous report, the anxious state is reflected by the reversed correlation between the neuronal activities and open arm exploration time^48^, we therefore performed *in vivo* calcium signal recordings, and further confirmed that activation of accumbal PV neurons is well correlated to avoidance of anxiogenic EPM open arms (Fig. 3n).

Indeed, other observations in our present study also support this hypothesis. First, we used *in vivo* single-unit recordings to confirm the relationship between anxiogenic stimuli and the activity of accumbal PV neurons. Among our recorded neurons, a type of fast spiking (FS) unit showed preferential activity in the anxiety-inducing environment (open arms, Fig. 3c-d, Fig. S2a). Because most fast-spiking neurons within the NAc have previously been identified as PV neurons, it confirms that accumbal PV neurons were engaged in the anxiogenic-related behavior. Second, our results (Fig. 3k) were consistent with a recent study showing that excitation of D2R cells in the NAc was necessary for normal, innate risk-avoidance^16^; however, we further demonstrated that PV neurons fired earlier than D2R cells (Fig. 3l-m), which implies that the accumbal PV neurons are informed of the risk and then modulate the activity of nearby cells that, in turn, initiate avoidance coping behavior. Further study is needed to investigate the accumbal neuronal firing dynamics in orchestrating risk avoidance in response to anxiogenic stimuli. Third, *in vivo* genetic manipulation of PV-neuronal activity in both healthy and chronic stress models further highlighted their importance in encoding anxiety-related behavior (Fig. 2i-k and Fig. 4). Taken together, these findings significantly extend previous observations of NAc in anxiety-related information processing; this is the first study that has begun to reveal real-time PV activity in the accumbens during the encoding of anxiety-related behavior in free exploration of aversive spaces without prior training and suggests that the underlying neuronal mechanism has been evolutionarily programmed.

Previous studies have showed that the NAc receives intermingled glutamatergic and dopaminergic inputs from a variety of forebrain regions, including the amygdala^23^, hippocampus^22^, thalamus^21, 33^, ventral tegmental area^19^ and the prefrontal cortex^34^. Using Cre-dependent, rabies-virus-based whole brain monosynaptic tracing strategy and electrophysiological recordings from brain slices, we demonstrated that NAc PV cells were specifically innervated by the GABAergic afferents stemming from the adBNST (Fig. 5 and Fig. 6d). To our best knowledge, this is the first study to map novel neural circuitry specifically innervating NAc PV neurons. Although a recent study found that light-evoked activation of vHPC inputs to NAc resulted in NAc^PV^ cell regulation of cocaine-seeking behavior, a cell-specific monosynaptic tracing strategy was not used in their study to show the anatomic connection^22^. Therefore, we speculate that the PV response to vHPC afferent activation may be indirect.

A previous study has shown that anterior BNST-associated activity exerts anxiolytic influence on anxious states^49^. Several of our current findings are consistent with this conclusion: specific activation of these afferents from the adBNST resulted in a robust inhibition of accumbal PV activity (Fig. 6d) and reduced avoidance coping behavior in response to anxiogenic stimuli (Fig. 6i-j); when PV function was ablated, the previously observed reduced avoidance behavior was also abolished (Fig. 6k-n); PPR has been reported to reflect the state of presynaptic input at synapses^50^. In our chronic stress model, we found in the adBNST-- sNAc circuit, there was an increase in the PPR, which indicating that an impaired release of GABA onto sNAc^PV^ cells unpon light activation of these GABAergic terminals in the sNAc (Fig. 6o-q); meanwhile, elevating the activity of GABAergic inputs to the NAc^PV^ neurons rescued anxiety-related excessive avoidance behavior, representing an anxiolytic effect (Fig. 6r-t). Combine these findings, we summarised that adBNST sents GABAergic inputs to sNAc to control avoidance behaviour, which is mediated by sNAc^PV^ neurons.

PV activity has been implicated in contributing to the *theta* rhythms in the mPFC and hippocampus^25, 26^ and our results also showed a marked decrease in *theta* rhythm on stressed mice both in NAc and BNST (Fig. 1j-k); additionally, electrophysiological recordings from freely-moving mice confirmed consistently lower local *theta* power in healthy animals during exploring the anxiogenic open arms (unrelated to locomotion, Fig. S2c-e), suggesting a strong correlation between *theta* power changes and anxious states. We reason that accumbal PV neurons exhibited high excitability under anxious states and therefore, highly activated PV cells contribute to *theta* oscillations changes either in BNST or NAc. Further study is needed to determine whether PV neuronal activation in the accumbens is the main driver for the decrease in *theta* oscillation within these two structures (Fig. 1j-k), although in the present study the increase in the coherence between FS spikes and LFP in *theta* range does suggest that this may be the case (Fig. S2f).

In conclusion, our results provide strong evidence for accumbal PV neurons driving anxiety-related avoidance coping behavior and may provide a new basis for the therapeutic purpose of pathological anxiety. Despite being a relative minor (∼4%) component of all NAc neurons^22^, this population has a robust anxiety-related behavioral effect. These accumbal PV cells are innervated by a long GABAergic-projecting input stemming from the adBNST. Anxiety represents a brain state: our study uncovered a new circuit mechanism, precisely defined by the neuronal types involved, by which the stress response brain region orchestrates the reward circuit component to exert direct effects on anxious states. Our findings may help to explain why anxiety and addiction are highly comorbid, although these two common psychiatric disorders engage emotion and reward circuits, respectively.

## Acknowledgements

This work was funded by the National Natural Science Foundation of China (31671116 J.T., 31761163005 J.T., 31800881 L.W. and 91132306 L.W.), the National Science Fund for Distinguished Young Scholars of NSFC (81425010 L.W.), the International Big Science Program Cultivating Project of CAS (172644KYS820170004 L.W.), the National Basic Research Program of China (973 program: 2015CB856402 J.T.), the External Cooperation Program of the Chinese Academy of Sciences (172644KYSB20160057 J.T.), the Youth Innovation Promotion Association of the Chinese Academy of Sciences (2017413 P.W.), the Guangdong Provincial Key S&T Program (2018B030336001 J.T.), Shenzhen Government Basic Research Grants (JCYJ20160429190927063 J.T., JCYJ20170413164535041 L.W.), Shenzhen Discipline Construction Project for Neurobiology DRCSM [2016]1379 (L.W.). We are gratefully for the comments and advice on the manuscript given by Prof. Liqun Luo. We thank Prof. Zilong Qiu for PV-Cre, D1R-Cre and D2R-Cre mice. We also thank Mr. Xu ZB and Mr. Liu BF for their help in transgenic mice husbandry and phenotyping. We are grateful to Mr. Liu XL and Ms. Li NN for their help in virus packaging.

## Author contributions

J.T. and L.W. conceived of this study. Q.X., X-Y.Z., L.X., B-F.W., and A-L. C. performed experiments. J.T., Q. X., X-Y. Z., P-F.W., Y-N.H., A-L. C. and J.W. analyzed data. Y.L. and F-Q.X. helped to design the experiments and provided suggestions. J.T. and Q.X. wrote the manuscript.

