## supplemental information for "A new GABAergic projection from the BNST onto accumbal parvalbumin neurons controls anxiety"

Figure Sup 1. Resting-state fMRI functional connectivity analysis.

Figure Sup 2. Accumbal theta power is coherent with fast spiking neuronal activitiy.

Figure Sup 3. NAc Ca^2+^ transients of different eYFP-tagged neuron populations in the EPM.

Figure Sup 4. *In vivo* recording of optogenetically-tagged NAc PV and MSN cells.

Figure Sup5. Negative control for retrograde tracing of the input neurons that send afferents to sNAc^PV^ neurons.

Figure Sup 6. *In situ* hybridization for identifying the neuronal types innervating downstream sNAc^PV^ neurons

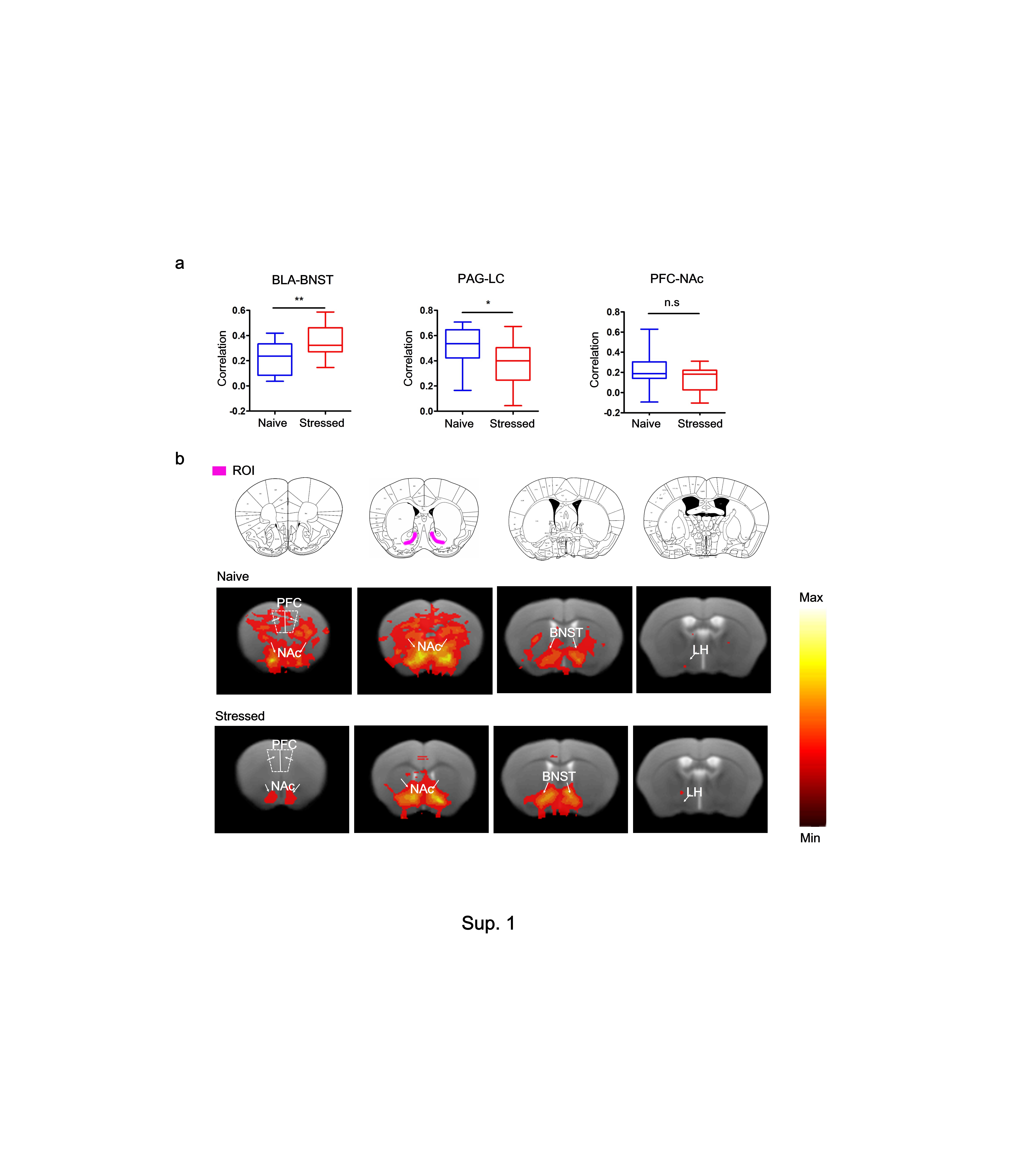

**Sup 1. Resting-state fMRI functional connectivity analysis**

(a) BOLD synchronization in BLA-BNST, PAG-LC and PFC-NAc in stressed and naive littermates (n = 5-7, unpaired *t* test, **P* < 0.05, ***P* < 0.01). (b) Heat maps showing a correlation of resting-state fMRI BOLD signal across brain regions viewed in coronal sections with a seed (ROI) in NAc.

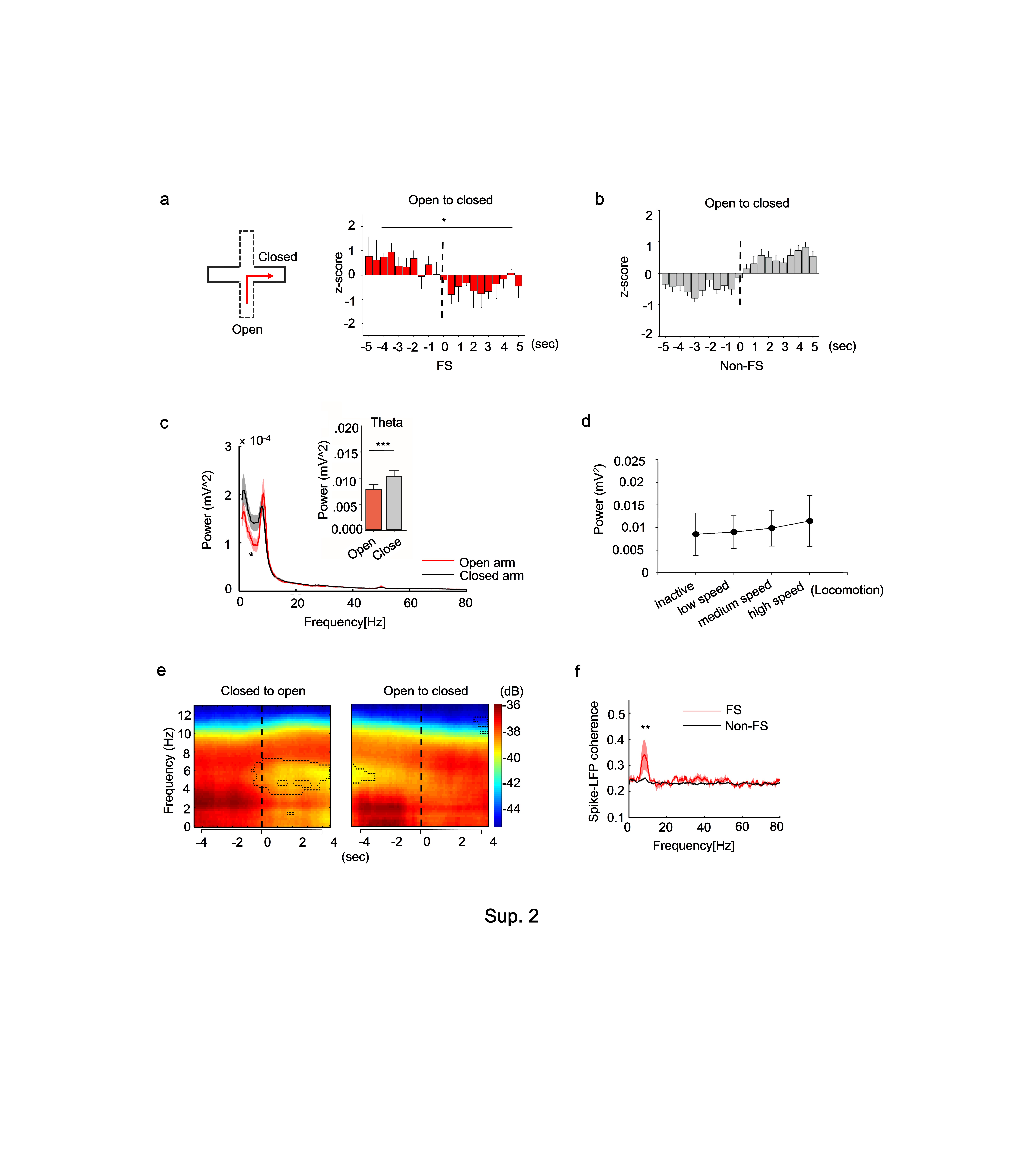

**Sup 2. Accumbal theta power is coherent with fast spiking neuronal activitiy**

(a) *Left,* horizontal and vertical arms represent closed and open arms, respectively; *right,* Z-scores of FS neurons during movement from the open to closed arms (nonparametric Kolmogorov-Smirnov test, n = 5, **P* < 0.05, see Methods). (b) There was no difference in individual Non-FS neuron firing rates from open to closed arms. (c) LFP within NAc shell in open and closed arms; *inset*, difference in the local theta activity between open and closed arms (Paired *t*-test, n = 21, *t* = -3.877 with 20 degrees of freedom, ****P* = 0.0009). (d) Correlation between locomotion and FS neuron LFP. (e) Average peri-event LFP spectrogram showing changes in the 4-8 Hz band during movement from the closed to open arms or from the open to closed arms. (f) Spike-LFP coherence at 4-8 Hz theta power of FS neurons and Non-FS neurons (Mann-Whitney rank sum test, n of FS = 5, n of Non-FS = 28, *t* = 141, ****P* = 0.005).

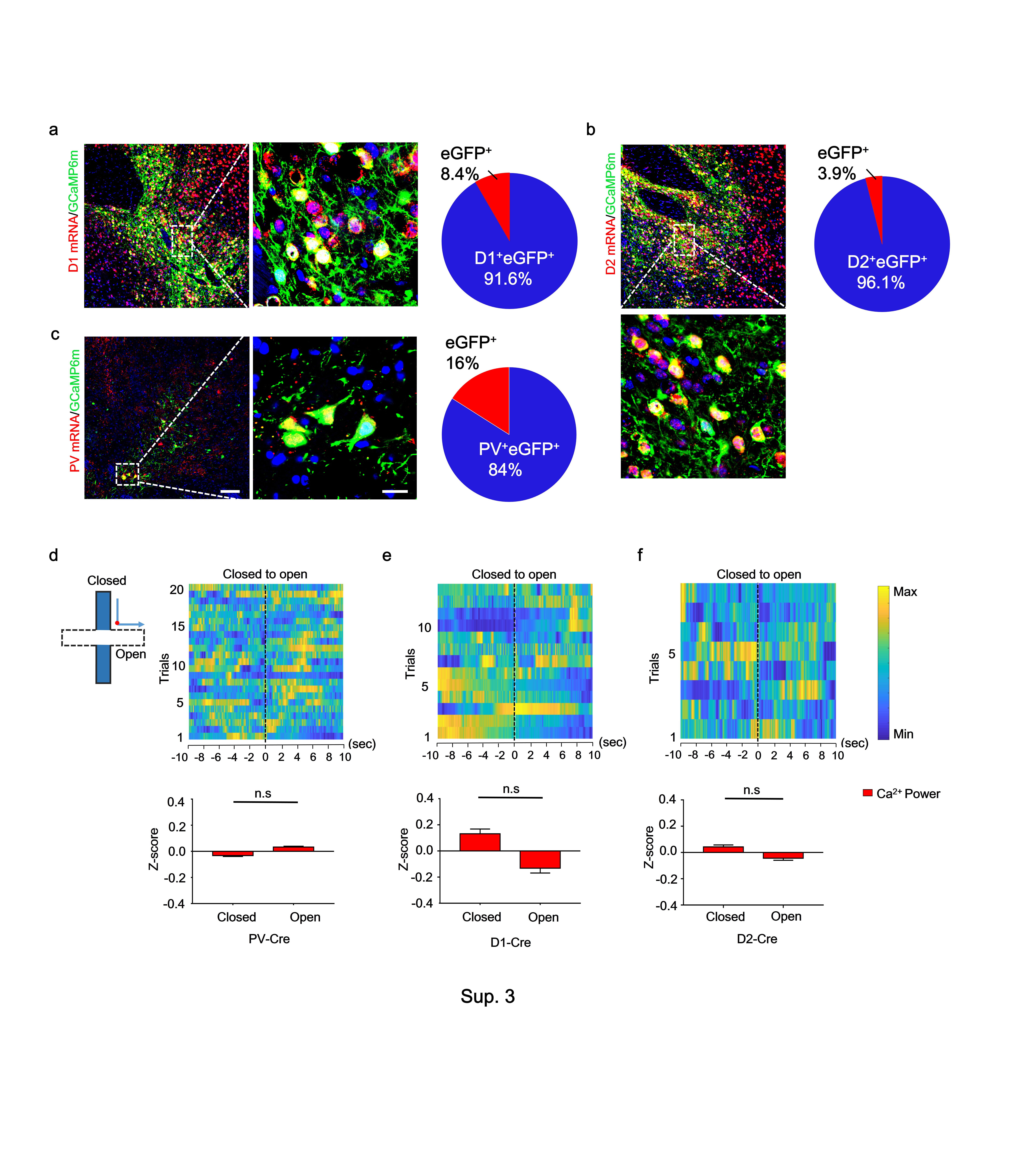

**Sup 3. NAc Ca^2+^ transients of different eYFP-tagged neuron populations in the EPM**

(a-c) Sections were co-stained with antibodies against eGFP (green) and riboprobes (red) for *D1R mRNA* (A, *left*)*, D2R mRNA* (b, *left top*) *and PV mRNA* (c, *left*); scale bar, 100 μm; enlarged view of the white box region showing cells co-expressing *D1R mRNA* (a, *middle)*, *D2R mRNA* (b, *left bottom)*, *PV mRNA* (c, *middle*) and GCaMP6m, respectively; scale bar, 10 μm, n = 9 slices from 3 mice per group; *right*, pie chart showing the respective co-expressed quantities of each mRNA type (*D1R mRNA* 91.6%*, D2R mRNA* 96.1%*, and PV mRNA* 84%). (d-f) *Top,* heatmaps of normalized NAc Ca^2+^ activity from different eYFP-tagged neuron populations in the EPM, binned by time (s) from the EPM crossing point (*inset*, red point). *Inset,* horizontal and vertical arms represent open and closed arms, respectively; *bottom*, normalized NAc Ca^2+^ transients of different eYFP-tagged neuron populations in the EPM open arms compared to closed arms (Wilcoxon test, n = 250).

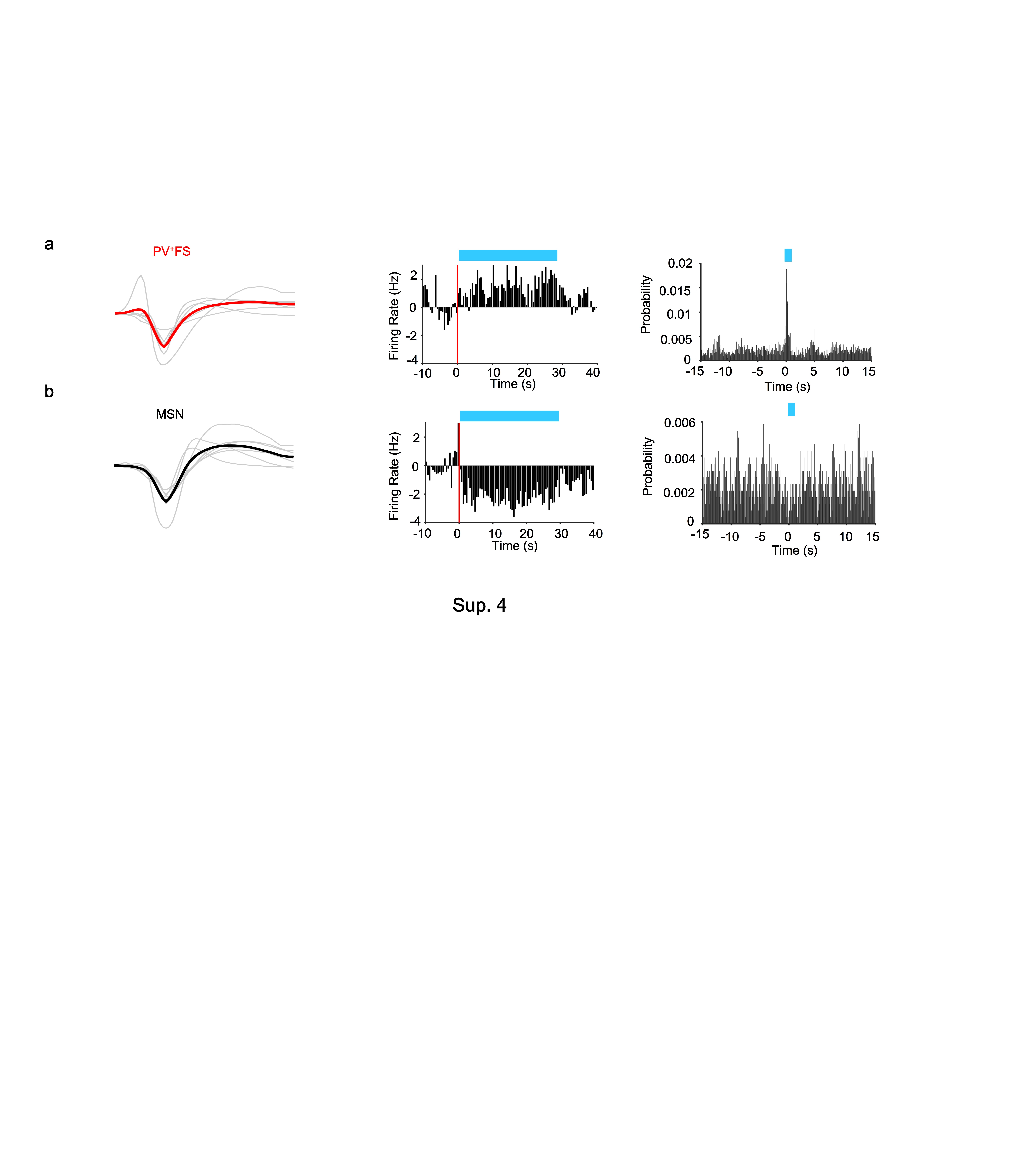

**Sup 4. *In vivo* recording of optogenetically-tagged NAc PV and MSN cells**

Examples of (a) an optogenetically-tagged NAc PV^+^ FS and (b) MSN recorded *in vivo*. From left to right, example spikes (gray traces) with superimposed mean waveforms (red/black traces) *(left*) in Light ON conditions, raster plots showing normalized short-latency light-driven spiking of FS units and decreased spiking of the MSN (*middle*), and their spike firing probabilities (*right*). The sharp peak at short latency identifies this FS unit as an optogenetically-tagged PV^+^ cell.

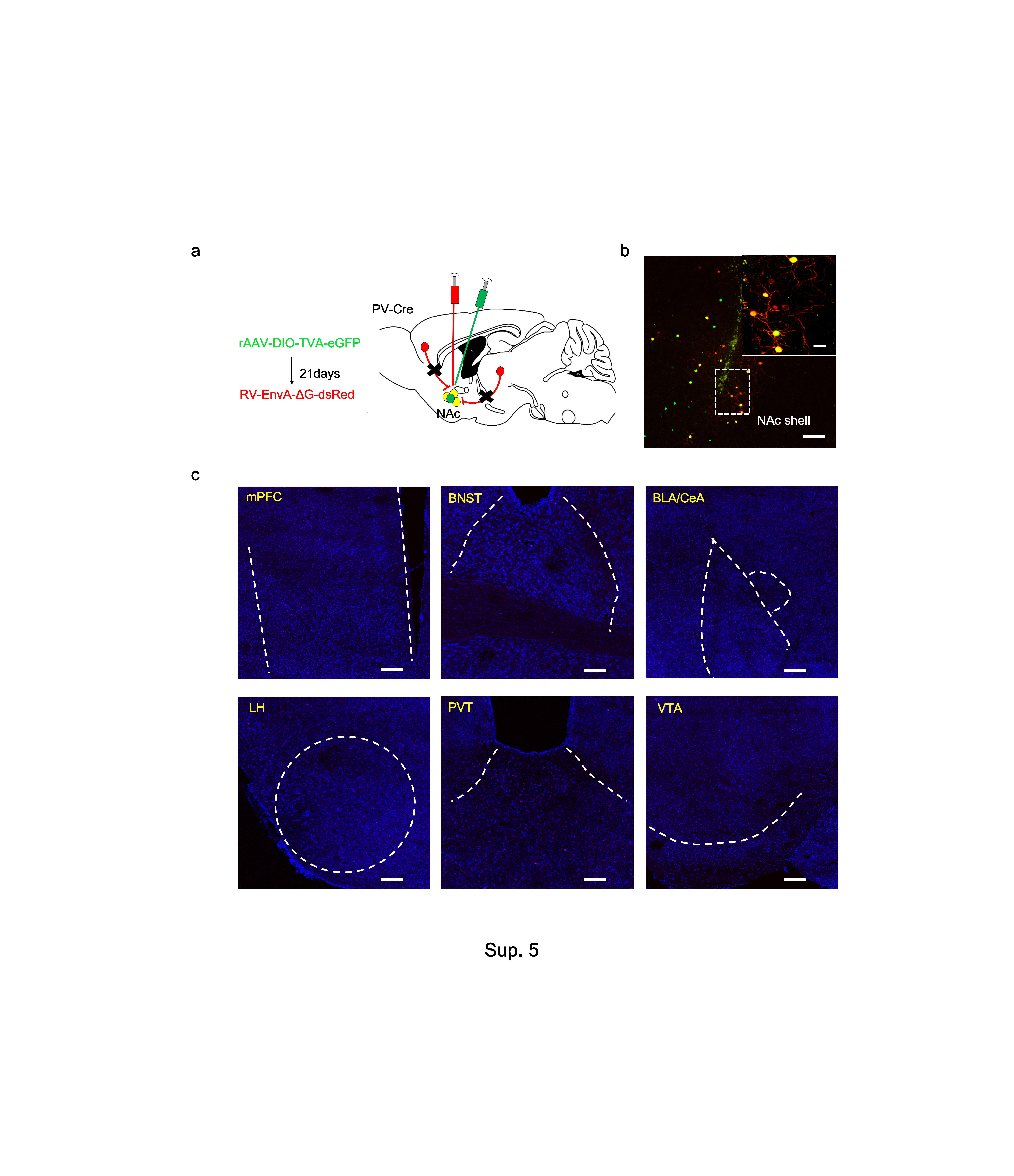

**Sup5. Negative control for retrograde tracing of the input neurons that send afferents to sNAc^PV^ neurons**

(a)Schematic showing injection of AAV-Ef1α-DIO-TVA-eGFP (AAV2/9) virus on day 1 and RV-EvnA-DsRed on day 21 into the sNAc of PV-Cre mice to retrogradely trace the input neurons (red) to NAc shell (yellow, starter neurons). (b) Fluorescence images of NAc region (coronal diagram) in PV-Cre mice (n = 4 mice), scale bar, 100 μm; *inset*, enlarged view of the white box region showing starter cells (yellow, expressing both eGFP and DsRed. Scale bar, 50 μm.) (c) No DsRed signals found in related brain regions.

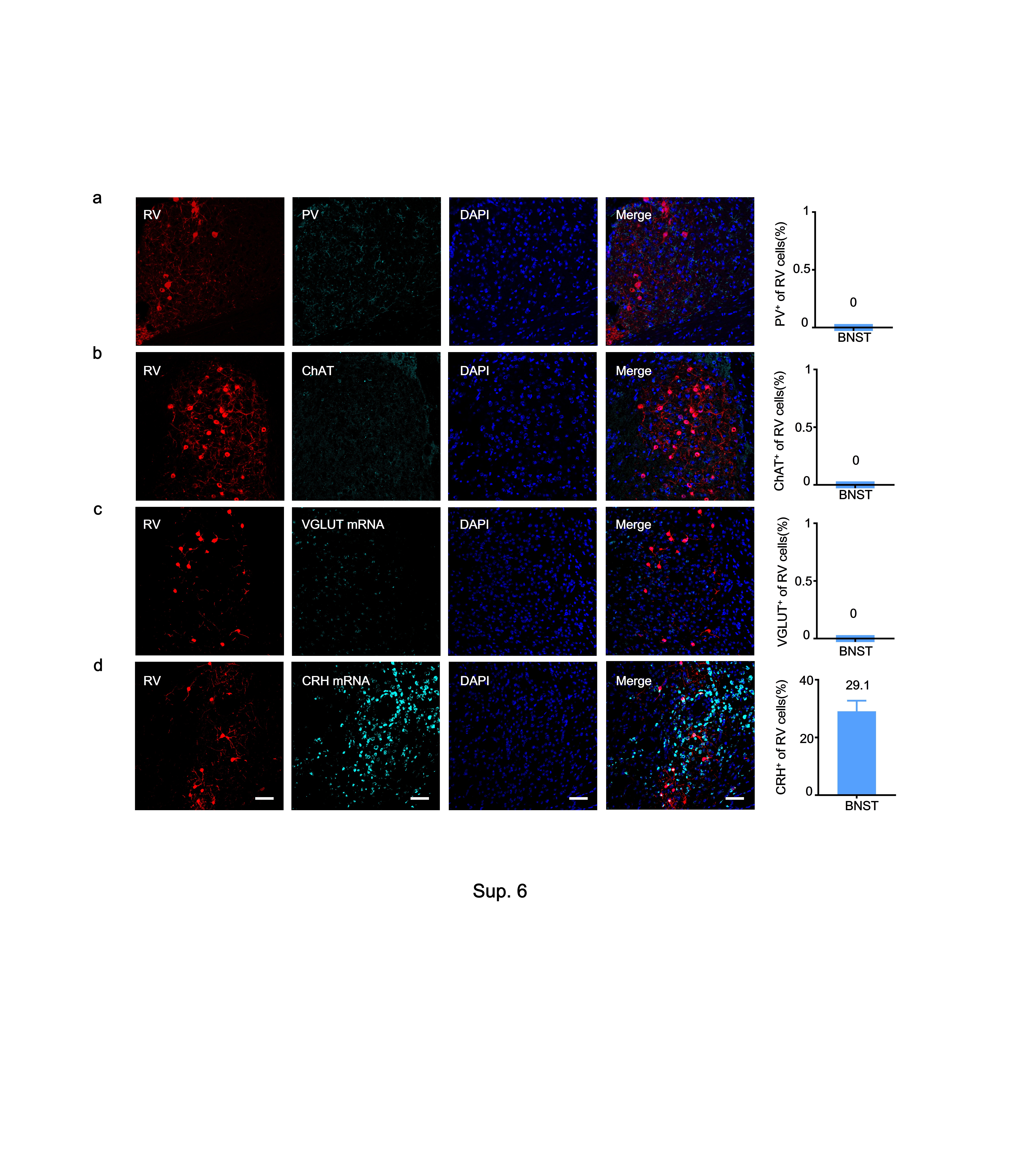

**Sup 6. *In situ* hybridization for identifying the neuronal types innervating downstream sNAc^PV^ neurons**

(a-c) *Left*, representative images showing no expression of PV, ChAT or VGLUT in the aBNST; *right,* quantitative bar showing the RV cells were neither expressing PV, ChAT nor VGLUT (n = 9 from 3 mice). (d) *Left*, representative images showing sparse co-expression of CRH and RV signals in the aBNST, scale bar, 50 μm; *right*, quantitative bar showing around 29.1% RV-cells were co-expression of CRH (n = 6 from 2 mice).
